# Antigen-aware TCR embeddings reveal conserved immune patterns for personalized immune monitoring

**DOI:** 10.64898/2026.10.09.757444

**Authors:** Mingyao Pan, Jing Hu, Yizhou Tracy Wang, Yuhao Tan, Hailong Hu, Ziqi Yang, Julia Fleming, Yuanyuan Pang, Bo Li

**Affiliations:** Department of Bioengineering, School of Engineering, University of Pennsylvania, Philadelphia, PA, USA; Center for Computational and Genomic Medicine, The Children’s Hospital of Philadelphia, Philadelphia, PA, USA; Department of Pathology and Laboratory Medicine, University of Pennsylvania, Philadelphia, PA, USA

## Abstract

T cell receptor (TCR) repertoires are highly individualized, while technical variation further complicates comparisons across individuals and cohorts. Here, we present TCRmeta, an antigen-aware framework that integrates population-scale repertoire relationships with experimentally defined antigen specificity to establish a shared space for repertoire analysis. TCRmeta predicted mutation-induced changes and captured antigen-related relationships in TCR specificity. Despite extensive sequence diversity, TCRmeta revealed a conserved repertoire landscape, enabling construction of a healthy reference for quantifying repertoire dispersal. Across cohorts, aging was associated with conserved V gene changes and progressive dispersal, with greater dispersal in CMV-positive individuals. Longitudinal analysis showed that repertoires progressively shifted toward the healthy reference following bone marrow transplantation or antiretroviral therapy. To directly compare repertoires in TCRmeta space, we developed Energy Shift, a pairwise measure robust to technical variation and HLA background while remaining sensitive to biological differences. Together, TCRmeta enables quantitative cross-individual and longitudinal immune monitoring.

**Teaser:** TCRmeta reveals conserved organization underlying highly individualized TCR repertoires and enables robust tracking of repertoire changes across immune states.

## Introduction

T cell receptors (TCRs) are highly diverse antigen-recognition receptors that enable T cells to recognize a broad array of antigens^1^. Exposure to different antigens drives the selection and expansion of antigen-specific T cell clones and shapes an individual’s TCR repertoire^2, 3, 4^. TCR repertoire composition therefore reflects an individual’s antigen experience and ongoing immune responses, making it a promising readout for longitudinal immune monitoring. However, the highly individualized nature of TCR repertoires complicates their clinical application^5^. The same antigen exposure or biological condition may expand different TCR clonotypes across individuals^6, 7^, making shared TCR signals difficult to identify across populations. This variability is further compounded by incomplete repertoire sampling. Any observed TCR repertoire represents only a small fraction of an individual’s full repertoire, and even samples collected from the same individual at the same time can contain largely different TCR clonotypes^5, 8^. Differences in sequencing depth and batch effects introduce additional technical variation^5, 9^. As a result, observed differences between TCR repertoires represent a mixture of changes in immune state and variation arising from individual differences, repertoire sampling, and technical factors. Disentangling biologically meaningful signals from these sources of variation remains a major challenge for reliable immune tracking from TCR repertoire data.

Multiple computational approaches have been developed to compare highly diverse TCR sequences. Exact clonotype matching remains widely used to identify public TCRs across repertoires^7^, while methods based on sequence similarity extend these comparisons to related but non-identical TCRs^10^. More recently, deep-learning approaches and protein language models have enabled TCRs to be represented in continuous embedding spaces, with applications ranging from TCR clustering to antigen-specificity prediction^11, 12^. These advances provide an opportunity to move beyond individual TCR sequences and analyze diverse TCRs within a unified representation space. In this study, we sought to establish a meta-level TCR representation in which TCRs from different individuals and cohorts can be directly compared within a common biological framework. Since antigen recognition is the primary function of TCRs, we reasoned that this common space should be organized in antigen-related structures, positioning TCRs that recognize the same or related antigens closer together even when their sequences differ.

Here, we developed TCRmeta, an antigen-aware representation framework designed to establish this meta-level space. TCRmeta was trained using repertoire data from 4,299 individuals^13, 14^ to identify groups of related TCR sequences and benchmarked with existing TCR representation methods^6, 10, 11, 15, 16, 17, 18, 19^. TCRmeta captured mutation-induced functional changes across multiple deep mutational scanning (DMS) datasets and effectively identified antigen-related TCRs across independent datasets. At the repertoire level, TCRmeta revealed highly conserved repertoire architecture across individuals despite extensive sequence-level diversity. Large-scale analysis of young and aged individuals revealed progressive ‘flattening’ of the TCR distribution with age in the TCRmeta embedding space, which can be partially reversed by bone marrow transplantation or therapeutic intervention. Finally, we confirmed that TCRmeta is ideal for comparative repertoire analysis as it is robust to sequencing depth, batch effects, and HLA background while remaining sensitive to biological signals. Specifically, it identified a cancer-specific shift in the embedding space using blood samples collected from pre-diagnostic ovarian cancer patients. Together, these findings reveal conserved antigen-aware organization beneath extensive TCR sequence diversity and provide a foundation for personalized immune monitoring using TCR repertoires.

## Results

### TCRmeta learns antigen-aware representations of TCR sequences

TCRmeta is a two-stage framework that integrates antigen-related information from large-scale TCR repertoires with experimentally annotated antigen specificity (Fig. 1A). The framework first learns TCR relationships from unlabeled repertoire data through pre-training and then refines the embedding space using known antigen specificity. TCRmeta is developed based on the rationale that similar TCRs within the same individual are more likely to have been activated and expanded in response to the same antigens^6, 20^. In other words, shared TCR similarity is a surrogate for shared antigen specificity.

**Fig. 1.**
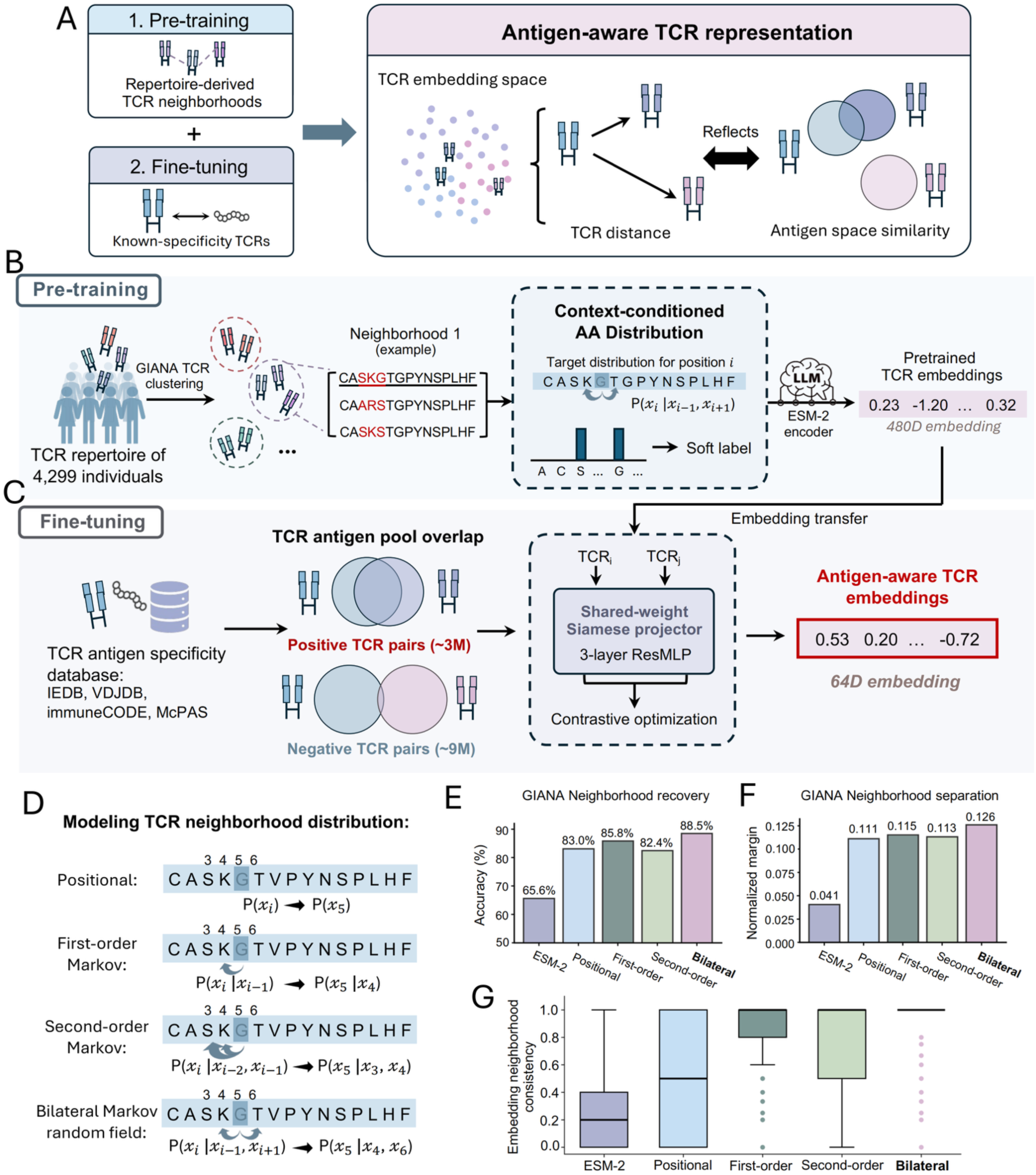
TCRmeta framework for learning antigen-aware TCR representations. **(A)** Overview of the TCRmeta framework, consisting of repertoire-based pre-training followed by fine-tuning with antigen-specific TCRs. **(B)** Pre-training framework. TCR neighborhoods were constructed from 4,299 TCR repertoires using GIANA clustering, and neighborhood-derived amino acid (AA) distributions were used as soft labels to pre-train the ESM-2 encoder. **(C)** Fine-tuning framework. TCRs with known antigen specificity from IEDB, VDJdb, immuneCODE, and McPAS-TCR were used to construct positive (3,007,593 pairs) and negative (9,790,950 pairs) TCR pairs for contrastive optimization, generating 64-dimensional antigen-aware embeddings. **(D)** Illustration of four strategies for modeling neighborhood AA distributions: positional, first-order Markov chain, second-order Markov chain, and bilateral Markov random field. **(E-F)** Comparison of different pre-training objectives in preserving GIANA-defined TCR neighborhoods. For each query TCR, positive TCRs from the same GIANA neighborhood were randomly selected together with control TCRs from outside that neighborhood. The control TCRs had the same Hamming distance and mutation positions relative to the query TCR as the positive TCRs. **(E)** Percentage of cases in which the positive TCR was closer to the query TCR than the control TCRs on average. **(F)** Mean normalized distance margin, calculated as (*d_control_ – d_positive_*)/(*d_control_ + d_positive_*); higher values indicate that positive TCRs are placed closer to the query than control TCRs. **(G)** Neighborhood consistency of ESM-2 and the four pre-training strategies in the TCR specificity datasets. For each model, TCR neighborhoods were defined based on distances in that model’s embedding space using the model-specific distance threshold (see Methods). Neighborhood consistency was calculated as the proportion of TCRs within the neighborhood that shared at least one antigen binder with the query TCR.

Based on this rationale, we used TCR sequence clusters from unlabeled repertoire data as a source of antigen-related information in the pre-training step. We collected 4,299 TCR repertoires^13, 14^ and applied GIANA clustering^10^ independently to each repertoire to identify clusters of similar TCRs. For each TCR, we then define its ‘neighborhood’ as the union set of all the GIANA clusters containing that TCR across samples in the training data. We retained TCRs with at least 10 distinct neighbors, resulting in 134,664 TCRs for pre-training. The AA composition of each neighborhood was summarized as position-specific probability distributions and used as soft labels to pre-train the ESM-2 encoder^19^ (Fig. 1B). By learning the AA alternatives observed among neighboring TCRs, we anticipate the model to learn which sequence substitutions are more likely to preserve related antigen specificity.

We next incorporated known paired TCR/antigen information through fine-tuning (Fig. 1C). Our goal was not to predict the antigen of each TCR, but to use antigen annotations to refine the geometry of the embedding space so that TCRs with related antigen specificity are closer by Euclidean distance. While pre-training learned antigen-related information from TCRs with similar sequences, this step allows limited expansion to learn how dissimilar TCRs preserve antigen specificity. We froze parameters in the pre-trained encoder layers and added a lightweight residual multilayer perceptron (ResMLP) for fine-tuning. The input data comprised pairs of TCRs, which were defined to account for TCR cross-reactivity, as a single TCR can bind to multiple peptide-major histocompatibility complex (pMHC)^21^. Specifically, we constructed TCR pairs from TCR/pMHC databases (IEDB^22^, VDJdb^23^, immuneCODE^14^, and McPAS-TCR^24^) using Fisher’s exact test and identified pairs with enriched or depleted overlap of antigen recognition (Extended Data Fig. 1A; see Methods). This generated 3,007,593 positive (enriched) and 9,790,950 negative (depleted) TCR pairs. Contrastive learning with these pairs produced the final 64-dimensional TCRmeta embedding for the downstream analyses.

A central step in TCRmeta is to distill the context-dependency for each AA substitution in a TCR neighborhood, which is the basis of CDR3 motifs in antigen recognition^18^. Hence, we evaluated different ways to extract this information. Specifically, we tested four strategies to model the probability distribution of AAs at each CDR3 position during pre-training, referred to here as the “TCR neighborhood distribution”: positional model, first-order and second-order Markov chain, and bilateral Markov random field model (Fig. 1D; Extended Data Fig. 1B; see Methods). We first asked whether these different pre-training strategies enabled the models to place TCRs from the same neighborhood (defined by GIANA clustering) closer than those outside the neighborhood. This was evaluated in an independent cohort^7^, with TCRs overlapping with the training set removed before redefining TCR neighborhoods. For each query TCR, positive TCRs were randomly selected from the same neighborhood. Control TCRs were selected from outside the neighborhood and they had the same Hamming distance and mutation positions from the query as the positive TCRs. This design allowed us to test which model(s) distinguish TCRs with identical sequence mismatches and learned the clustering context. All four pre-training strategies improved neighborhood recovery compared with the ESM-2 baseline. Among them, the bilateral model performed best, achieving an accuracy of 88.5% (Fig. 1E). It also showed the largest normalized distance margin, indicating greater separation between positive and control TCRs relative to the query TCR (Fig. 1F). Based on these results, we selected the bilateral model as the final pre-training strategy for TCRmeta.

We next compared the different pre-training models using neighborhood consistency, defined as the proportion of nearby TCRs in the embedding space that shared at least one antigen binder with the query TCR (Methods). All four strategies improved neighborhood consistency compared with the original ESM-2 representation, while the bilateral Markov random field model showed the highest consistency (Fig. 1G). This is likely due to the fact that the bilateral model integrated both flanking regions and obtained a more accurate representation of the TCR neighborhood. Together, these results indicate that the bilateral model better preserves clustering-defined TCR relationships and antigen-related local structure in the embedding space.

### TCRmeta captures mutation-induced functional changes and antigen-related TCR relationships

We next asked whether pre-trained TCRmeta embedding could capture the functional effects of subtle TCR sequence variation. To test this, we evaluated the pre-trained model using published deep mutational scanning (DMS) datasets. DMS systematically introduces individual AA substitutions into a TCR and experimentally measures how each mutation affects TCR function (Fig. 2A). Seven TCRs from five published DMS studies^25, 26, 27, 28, 29^ were included, paired with 12 experimental readouts of TCR-pMHC binding or T cell activation across tumor and viral antigens (Fig. 2B; Extended Data Fig. 2A). Pre-trained TCRmeta was compared with four existing deep-learning methods trained on TCR sequences: CDR3vec^16^, TCR-VALID^15^, TCR-BERT^11^, and SCEPTR^17^. For each TCR mutant, we calculated its embedding distance from the wild-type TCR and compared this distance with the experimentally measured fitness score, with higher fitness scores indicating larger changes in TCR function relative to the wild type. If the embedding captures mutation-induced impacts in TCR specificity, the embedding distance is expected to change consistently with the experimentally measured fitness score.

**Fig. 2.**
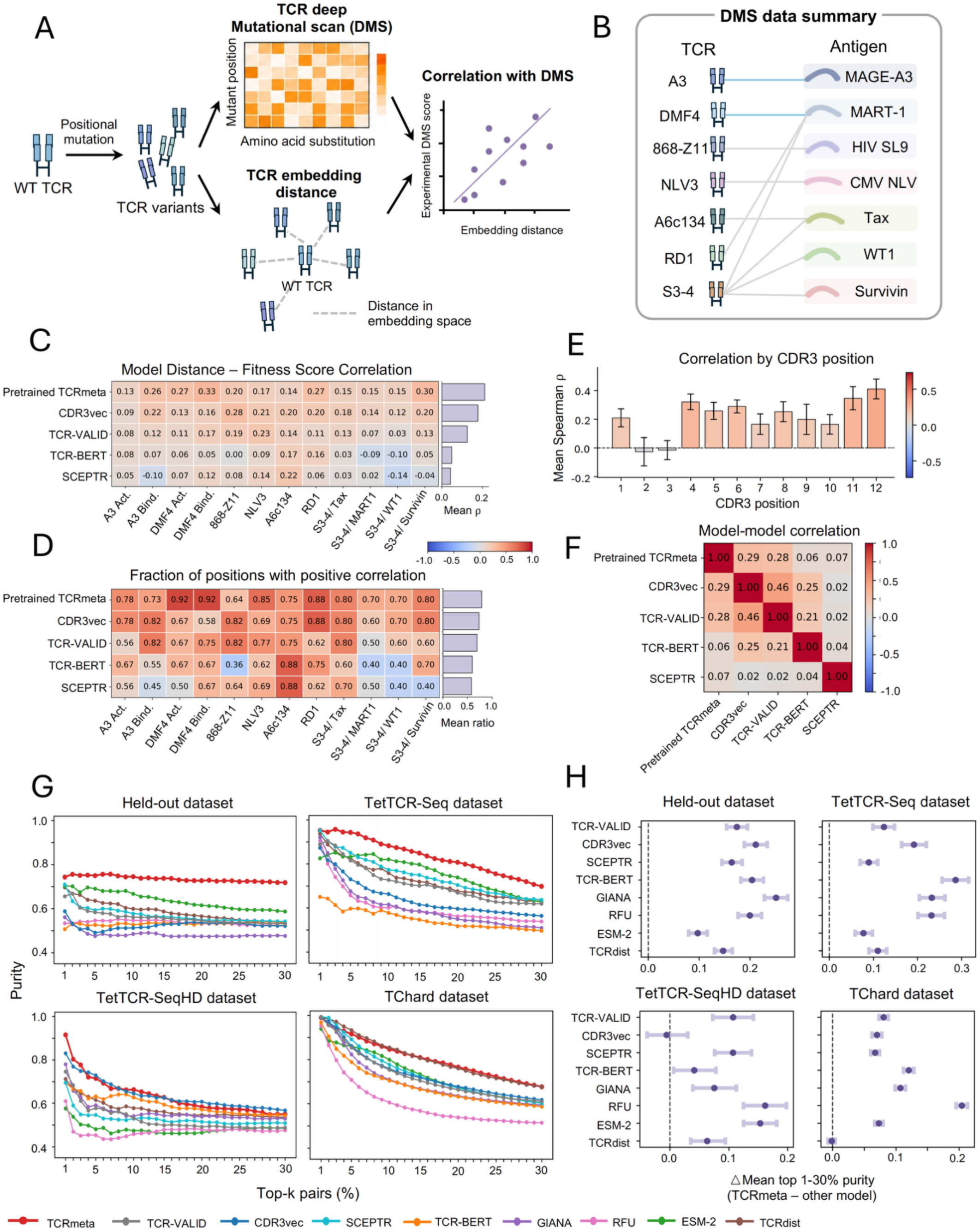
TCRmeta captures mutation-induced functional changes and antigen-related TCR relationships. **(A)** Schematic of the deep mutational scanning (DMS) analysis. Distances between wild-type TCRs and their single-AA variants in the embedding space were compared with experimental fitness scores. **(B)** Summary of the seven TCRs included in the DMS benchmark and their corresponding antigens. The datasets comprise 12 experimental readouts of TCR-pMHC binding or T cell activation from five published DMS studies. Blue lines indicate TCRs with both TCR-pMHC binding and T cell activation readouts, whereas gray lines indicate TCRs with binding readouts only. **(C)** Spearman correlations between mutant-WT embedding distance and experimental fitness score across the 12 DMS readouts. Values were calculated separately by CDR3β position and averaged across positions for each readout. Mean correlations across readouts are shown on the right. **(D)** Fraction of CDR3β positions showing a positive correlation between embedding distance and experimental fitness score for each DMS readout. **(E)** Mean Spearman correlation between pre-trained TCRmeta distance and experimental fitness score at each CDR3β position across DMS readouts. Error bars indicate standard error of the mean (SEM). **(F)** Pairwise correlations of TCR distance percentile between the five TCR embedding models. Distances were converted to percentile ranks within each DMS readout and CDR3β position before calculating Spearman correlations between models. Values represent the mean correlation across DMS readouts. **(G)** Purity of antigen-related TCR pairs among the top 1-30% nearest pairs in the held-out, TetTCR-Seq, TetTCR-SeqHD, and TChard datasets. TCR pairs were ranked by embedding distance, and purity was calculated as the fraction of positive antigen-related pairs within the indicated top percentage. **(H)** Difference in mean top 1-30% purity between TCRmeta and each comparison method. Points indicate the median difference across 5,000 bootstrap resamples of TCR pairs, and error bars indicate the 95% percentile interval. Positive values indicate higher purity for TCRmeta.

Indeed, TCRmeta is positively correlated with all 12 readouts, showing the highest mean correlations among the five methods (mean ρ = 0.21; Fig. 2C). In addition, the majority of CDR3β positions showed positive distance-fitness correlations, with proportions ranging from 0.64 to 0.92 across the 12 readouts (mean ratio = 0.79; Fig. 2D). For each CDR3β, mutation effects at positions 2 and 3 were more difficult to predict, whereas TCRmeta showed higher predictive power for positions 4-12 (Fig. 2E; Extended Data Fig. 2B). We also compared the five different TCR embeddings to assess the similarity of the information captured by each method. Interestingly, embeddings from the five methods were largely dissimilar, with correlations between pre-trained TCRmeta and the other methods ranging from 0.06 to 0.29 (Fig. 2F; Extended Data Fig. 3). These results suggest that different training strategies learn distinct signals from TCR sequences.

The DMS analysis showed that pre-training captured functional differences among closely related TCR sequences. Building on these local relationships, fine-tuning incorporated experimentally defined antigen relationships to connect dissimilar TCRs and establish a broader antigen-aware representation. We therefore asked whether the fine-tuned TCRmeta embedding could identify antigen-related TCRs beyond local sequence similarity. We performed evaluation using four unseen TCR/pMHC datasets: a held-out dataset, TetTCR-Seq^30^, TetTCR-SeqHD^31^, and TChard^32^. TCRs overlapping with the training data were excluded. Positive and negative TCR pairs were defined using the same enrichment approach as described in the fine-tuning step, resulting in 14,720, 6,300, 5,812, and 68,914 pairs, respectively (see Methods). For a more comprehensive evaluation, we expanded the benchmark to eight methods (TCR-VALID^15^, CDR3vec^16^, SCEPTR^17^, TCR-BERT^11^, GIANA^10^, RFU^18^, ESM-2^19^, and TCRdist^6^) spanning different TCR representation and distance strategies. Since only a small fraction of TCRs is expected to share antigens with a given TCR, we focused on the nearest TCR pairs, for which distance is most informative. Result showed that antigen-related TCRs were enriched among the nearest pairs in the TCRmeta space across all four datasets, which is statistically significant based on permutation test (Extended Data Fig. 4). TCRmeta also consistently showed the highest overall antigen-related pair purity across the nearest 30% of pairs compared with the other methods (Fig. 2G). We further quantified these differences using bootstrap resampling, calculating the difference in mean top 1% to 30% purity between TCRmeta and each comparison method (Fig. 2H). Positive values indicate higher purity for TCRmeta. Most comparisons showed positive differences across the four datasets, further supporting the strong predictability of TCRmeta for unseen, dissimilar TCRs across different benchmarks. Together, these results show that TCRmeta captures antigen-related relationships across diverse TCR sequences, providing a practical basis to model TCR repertoires in an antigen-aware embedding space.

### TCRmeta reveals a conserved organization of TCR repertoires across individuals

We applied TCRmeta to visualize TCR repertoires from diverse biological contexts, including healthy donors^14, 33^, type 1 diabetes (T1D)^33^, lung cancer^34^, recombination-activating genes (RAG) mutations^35^, thymocytes^36^ and TCR-like peptides as control (Fig. 3A). TCRs with different β-Variable (V) gene families showed clear separation in the embedding space, which is due to the V gene constraint in the pre-training step.

**Fig. 3.**
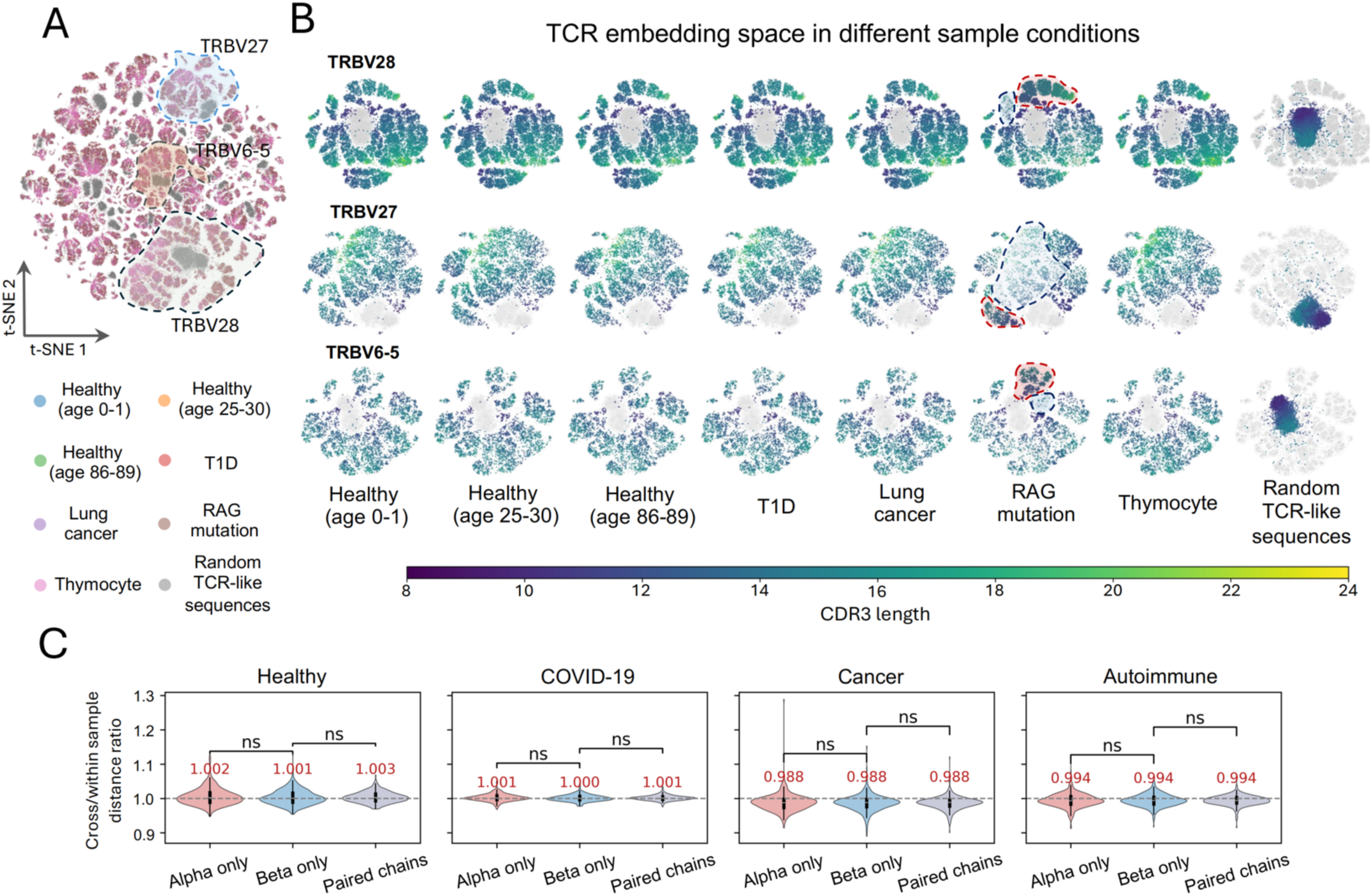
Conserved organization of the TCRmeta embedding space across repertoires. **(A)** t-distributed stochastic neighbor embedding (t-SNE) visualization of TCRmeta embeddings from healthy individuals of different ages, individuals with type 1 diabetes (T1D), lung cancer, or recombination-activating gene (RAG) mutations, and thymocytes, together with randomly generated TCR-like sequences as a negative control. Regions corresponding to the three most frequent TRBV genes (TRBV28, TRBV27, and TRBV6-5) are outlined. **(B)** TCRmeta embedding space shown separately for the three most frequent TRBV genes (TRBV28, TRBV27, and TRBV6-5). For each sample, TCRs are colored by CDR3 length, with TCRs pooled from all samples shown in gray as a reference background. Dashed outlines highlight regions enriched (red) or depleted (blue) in individuals with RAG mutations. **(C)** Comparison of repertoire organization using α-chain, β-chain, or paired αβ TCR embeddings in healthy, COVID-19, cancer, and autoimmune samples from the Observed TCR Space database. Within each TRAV-TRBV group, the cross/within-sample ratio was calculated as the mean distance between TCRs from different individuals divided by the mean distance between TCRs from the same individual. P values were calculated using two-sided Mann-Whitney U tests. ns, not significant.

To eliminate V gene-related batches, we focused on the three most frequent V genes, TRBV28, TRBV27, and TRBV6-5 (Fig. 3A-B). For each V gene, repertoires from different individuals or biological conditions showed a highly conserved distribution. This organization was also observed in thymocytes (Fig. 3B), suggesting that TCR repertoire structure is already established during early development of adaptive immunity. To examine whether V(D)J recombination is a cause for this conservation, we analyzed individuals carrying germline RAG mutations^35^ that directly disrupted the TCR generation process. Expectedly, repertoires from these patients showed clearly altered regions in the embedding space compared to healthy controls (Fig. 3B). We also examined randomly generated TCR-like sequences with matched length, C- and N-terminal CDR3 motifs. These non-biological sequences occupied a very different region of the TCRmeta space from biological TCRs (Fig. 3B). These results suggest that TCR generation contributes to the overall organization of TCR repertoires in TCRmeta space. Except for conditions that substantially disrupt TCR generation, TCR repertoire structure was largely conserved across individuals regardless of age or disease status.

We next asked if the observed conservation applies to both TCRα and TCRβ chains. We analyzed paired TCRαβ sequences from the Observed TCR Space (OTS) dataset^37^, including healthy individuals and patients with COVID-19, cancer, or autoimmune disease (Fig. 3C). To control for V gene differences, TCRs were grouped by their TRAV-TRBV combinations. Within each group, we calculated the ratio of distances between TCRs from different individuals to those from the same individual. Indeed, we observed ratios close to 1 for α, β, and paired αβ representations across all conditions (Fig. 3C), indicating that TCRs from different individuals were well mixed in the embedding space. Importantly, use of both α and β chains did not increase the separation between individuals (Fig. 3C), suggesting that the conserved organization observed using TCRβ alone is not due to missing α-chain information, but reflects a general feature of TCR repertoires. This also supports the use of TCRβ alone for capturing this repertoire-level organization. Together, these results reveal a conserved organization of human TCR repertoires despite substantial differences in their individual TCR sequences.

### TCRmeta reveals age-associated changes in TCR repertoire organization

The above conserved repertoire structure across individuals allowed us to construct a common reference in the TCRmeta space to study age-related repertoire-level changes. We used the Gittelman cohort, a large population-based study from Vo’, Italy^13^. The cohort included individuals spanning a broad age range. We selected 500 healthy individuals aged 0-30 years (4,348,248 TCRs) to construct a healthy immune reference map (Fig. 4A). For any new repertoires to be analyzed, their TCRs can be encoded by TCRmeta and projected onto this map. Their distribution can then be compared to the TCRs in the reference (Fig. 4A). To ensure sufficient TCR representation, we included 35 V genes with an average repertoire frequency above 1% (Extended Data Fig. 5A). We then used the remaining samples in the Gittelman cohort to examine how TCR repertoire organization changes with age. We randomly selected 100 individuals from each age group, except for the 21-30-year group, for which only 10 individuals were included because the remaining samples had been used to construct the young healthy reference. As an independent validation, we also analyzed the Emerson cohort^7^, a large U.S. TCR repertoire study published in 2017. The two cohorts differed in population, study background, and study period, allowing us to test whether the aging-associated pattern was reproducible across distinct cohorts.

**Fig. 4.**
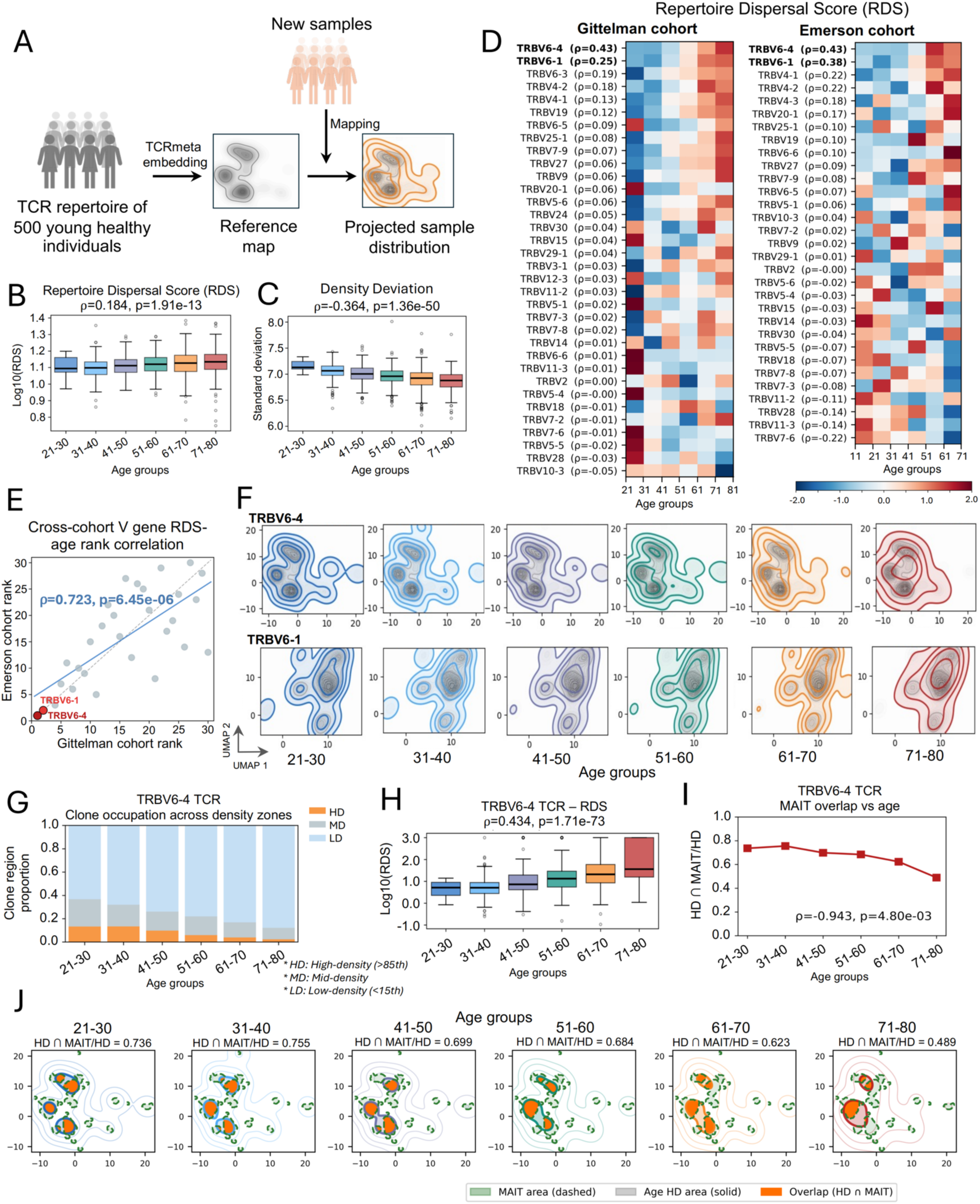
TCRmeta reveals age-associated changes in TCR repertoire organization. **(A)** Schematic of the Repertoire Dispersal Score (RDS). New repertoires were projected onto a reference map constructed from 500 healthy individuals, and RDS was calculated from the difference between the projected sample and reference distributions. **(B, C)** Overall **(B)** RDS and **(C)** density deviation across age groups. Spearman correlations with age are shown. **(D)** V gene-specific RDS across age groups in the Gittelman and Emerson cohorts. V genes are ranked by the Spearman correlation between RDS and age. **(E)** Correlation of V gene RDS rankings between the Gittelman and Emerson cohorts. **(F)** Uniform Manifold Approximation and Projection (UMAP) visualization of TRBV6-4 and TRBV6-1 TCR distributions across age groups in the Gittelman cohort. For each age group, the colored density shows the distribution of TCRs from that age group, while the gray background shows the distribution of the young healthy reference. **(G)** Fraction of TRBV6-4 clonotypes occupying high-density (HD), medium-density (MD), and low-density (LD) regions across age groups, defined by reference-density percentiles (>85th, 15th-85th, and <15th percentile, respectively). **(H)** TRBV6-4 RDS across age groups. Spearman correlation with age is shown. **(I)** Overlap between the high-density (HD) region of TRBV6-4 and the mucosal-associated invariant T (MAIT) cell area across age groups. **(J)** TRBV6-4 density distributions showing the HD region, MAIT area, and their overlap across age groups. Values above each panel indicate the fraction of the HD region overlapping the MAIT region.

We introduced the Repertoire Dispersal Score (RDS) to measure the distributional differences between a given repertoire sample and the reference. For each TCR, we measured its local density in the reference distribution. A TCR will then be classified as ‘high-density’ if its local density is above the 85th percentile of the reference density, or ‘low-density’ if below the 15th percentile. RDS was then calculated as the ratio of TCR occupancy in low-density regions to that in high-density regions (see Methods). A higher RDS indicates that more TCRs are distributed toward regions that are less densely occupied in the reference map. RDS increased significantly with age in the Gittelman cohort (ρ = 0.184, *P* value= 1.91 × 10^-13^; Fig. 4B), suggesting that TCR repertoires consistently shift from regions commonly occupied in the young healthy reference toward less frequently occupied regions in the TCRmeta space. We also calculated density deviation as the standard deviation of the density values across TCRs within each repertoire. Density deviation decreased with age (ρ = −0.364, *P value=* 1.36 × 10^-50^; Fig. 4C), indicating that compared to young donors, aged repertoires are more dispersed in the embedding space, with fewer extremities in the probability distribution. These trends were reproduced in the Emerson cohort, where RDS increased with age (ρ = 0.158, *P value=* 3.53 × 10^-2^; Extended Data Fig. 6A) and density deviation decreased (ρ = −0.377, *P value=* 2.07 × 10^-7^; Extended Data Fig. 6B).

We then investigated the relationship between RDS and age stratified by V gene. The correlations with age varied considerably across V genes (Fig. 4D). TRBV6-4 and TRBV6-1 showed the strongest positive correlations, and several members of the TRBV4 family also ranked in the top. We then examined whether the same V gene-specific pattern was present in the Emerson cohort. Despite the differences between the two cohorts, the V gene rankings were highly consistent, with TRBV6-4 and TRBV6-1 being the top two, followed by TRBV4 family members (Fig. 4D). Overall, the V gene rankings were strongly correlated between the two cohorts (ρ = 0.723, *P value=* 6.45 × 10^-6^; Fig. 4E), supporting a reproducible V gene-specific pattern of repertoire changes with aging.

We next visualized these changes in the TCRmeta space, focusing on TRBV6-4 and TRBV6-1. In the Gittelman cohort, repertoires from independent younger donors closely followed the shape of the reference map and showed sharp local features in distribution (Fig. 4F). With increasing age, the repertoire distribution progressively shifted away from the reference, and these fine-scale features became less pronounced as the overall landscape appeared smoother. These spatial changes were consistent with the increase in RDS and decrease in density deviation observed with age (Fig. 4B-C). A similar age-associated change in repertoire shape was observed in the Emerson cohort (Extended Data Fig. 6D). One possible explanation is that some of the fine-scale structures in younger repertoires are formed by public clones shared across individuals. Over time, the accumulation of individual-specific antigen exposures, clonal expansions, and other repertoire remodeling may make TCR occupancy increasingly individualized, gradually weakening these shared fine-scale structures at the population level.

We then sought to understand why some V genes, for example, TRBV6-4, showed consistently stronger associations with age. Specifically, the fraction of TRBV6-4 TCRs in high-density regions decreased with age, whereas the fraction in low-density regions has an opposite trend (Fig. 4G). TRBV6-4 RDS showed a strong positive association with age in the Gittelman cohort (ρ = 0.434, *P value=* 1.71 × 10^-73^; Fig. 4H), and a similar association was observed in the Emerson cohort (ρ = 0.428, *P value=* 2.45 × 10^-9^; Extended Data Fig. 6C). We hypothesized that the strong age association of TRBV6-4 TCRs might be related to mucosal-associated invariant T (MAIT) cells. This is because TRBV6-4 is preferentially used by MAIT cells, and circulating MAIT cell frequency has been reported to decline with age^5, 38^. To test this hypothesis, we collected MAIT TCR sequences from five published single-cell TCR-seq datasets^39, 40, 41, 42, 43^ and projected them onto the reference map to define MAIT area in the TCRmeta space (Fig. 4I-J). We then measured the fraction of the high-density region that overlapped with the MAIT area across age groups. For TRBV6-4, this overlap progressively decreased with age in the Gittelman cohort (Fig. 4I-J), with the same trend observed in the Emerson cohort (Extended Data Fig. 6E-F). These results suggest that the age-associated decline in MAIT cells is a likely cause to the age-related changes for TRBV6-4 TCRs. However, MAIT TCRs also frequently use TRBV6-1 and TRBV20-1, which showed little or no age-associated decrease in overlap in either cohort (Extended Data Fig. 5C-D; Extended Data Fig. 6E-F). Thus, changes in the MAIT population may partly explain the strong age association of TRBV6-4 but cannot fully account for the differences observed across V genes.

### Repertoire dispersal reflects immune state and response to clinical interventions

In the above section, we hypothesized that aging may cause a ‘flattened’ distribution due to prolonged and systematic antigenic response. We therefore asked whether chronic antigenic exposure can replicate a similar shift in the TCRmeta space. We first examined cytomegalovirus (CMV)^7^, a persistent viral infection with long-term effects on the T cell repertoire. In TRBV6-4 TCRs, RDS increased with age in both CMV-negative and CMV-positive individuals, but the association was stronger in the CMV-positive group (CMV-: ρ = 0.207, *P value=* 7.11 × 10^-5^; CMV+: ρ = 0.414, *P value=* 3.30 × 10^-14^; Fig. 5A). Across age groups, CMV-positive individuals generally showed higher RDS than CMV-negative individuals, particularly among older population (Fig. 5B). These results suggest that persistent CMV-associated immune stimulation may contribute to progressive repertoire dispersal.

**Fig. 5.**
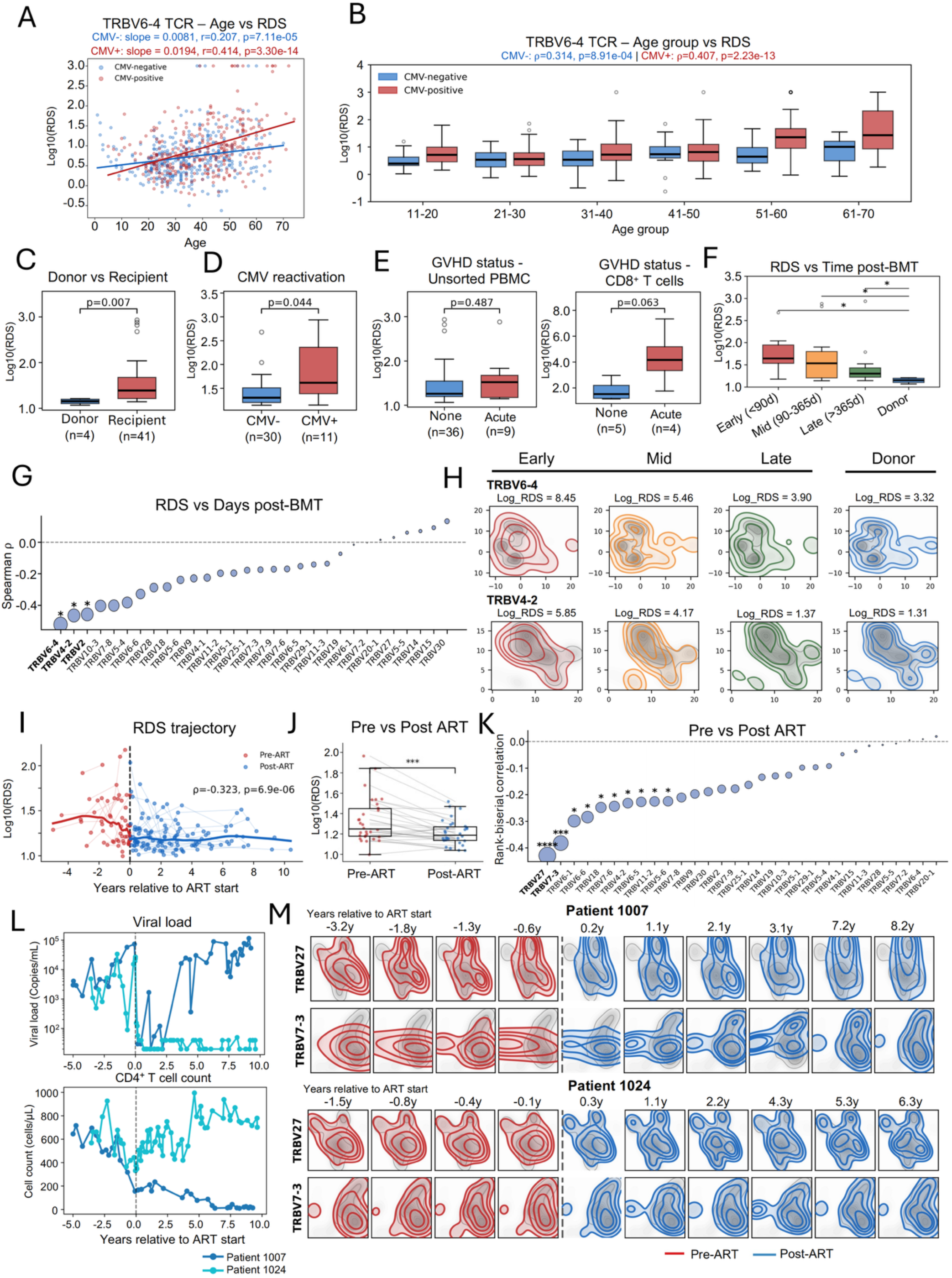
Repertoire dispersal reflects immune state and response to clinical interventions. **(A)** Association between age and TRBV6-4 Repertoire Dispersal Score (RDS) in CMV-negative and CMV-positive individuals. Lines show linear fits for each group. Spearman correlations are shown. **(B)** TRBV6-4 RDS across age groups stratified by CMV status. **(C)** RDS of donor and recipient repertoires following bone marrow transplantation (BMT). **(D)** RDS in recipients with or without CMV reactivation after BMT. **(E)** RDS according to graft-versus-host disease (GVHD) status in unsorted peripheral blood mononuclear cells (PBMCs) and CD8^+^ T cell populations. **(F)** RDS at early (<90 days), mid (90-365 days), and late (>365 days) time points after BMT compared with donor repertoires. **(G)** Spearman correlations between V-gene specific RDS and days post-BMT. V genes are ranked by correlation strength, and point size represents -log10(adjusted *P* value). **(H)** Density distributions of TRBV6-4 and TRBV4-2 at early, mid, and late time points after BMT and in donors. For each group, the colored density shows the distribution of TCRs from that time post-BMT group, while the gray background shows the distribution of the young healthy reference. **(I)** RDS trajectories before and after antiretroviral therapy (ART) initiation. The dashed line indicates ART initiation. Spearman correlation between RDS and time after ART initiation is shown. **(J)** Paired comparison of RDS before and after ART. Statistical significance was assessed using a two-sided paired *t*-test. **(K)** Rank-biserial correlations between V-gene-specific RDS and ART status. V genes are ranked by correlation strength, and point size represents -log10(adjusted *P* value). **(L)** Longitudinal viral load and CD4^+^ T cell count for patient 1007 and patient 1024 relative to ART initiation. **(M)** Density distributions of TRBV27 and TRBV7-3 before and after ART in patients 1007 and 1024. *: P value < 0.05; **: P value < 0.01; ***: P value < 0.001; ****: P value < 0.0001.

We next applied TCRmeta to study repertoire changes in patients who received allogeneic bone marrow transplantation (BMT)^44^. Post-BMT recipients showed significantly higher RDS than donors (*P value=* 0.007; Fig. 5C), indicating that the reconstituted repertoires remained more dispersed. Consistently, recipients with CMV reactivation showed higher RDS than those without (*P value=* 0.044; Fig. 5D). In patients with acute graft-versus-host disease (GVHD), RDS did not differ by GVHD status in unsorted PBMCs (*P value=* 0.487), whereas CD8⁺ T cells showed a modest trend toward higher RDS in patients with acute GVHD (*P value=* 0.063; Fig. 5E), suggesting subset-specific changes. Interestingly, we observed a continued decrease of RDS post-BMT, with recipients at later time points approaching the levels observed in healthy donors (Fig. 5F). This trend was observed across multiple V genes, with TRBV6-4 and TRBV4-2 showing the strongest associations with post-BMT time (Fig. 5G). Notably, TRBV6-4 and TRBV4-2 were also among the top V genes linked to aging in the earlier analysis, suggesting that post-BMT changes may partly counter age-related repertoire remodeling. Early post-BMT repertoires showed substantial deviation from the healthy reference, whereas later repertoires gradually regained structural features resembling the reference and donor repertoires (Fig. 5H). These results collectively indicated partial TCR repertoire rejuvenation after BMT from younger donors and confirmed the utility of TCRmeta in tracking TCR repertoire changes in clinical samples.

We then analyzed longitudinal TCR data from patients with human immunodeficiency virus (HIV) before and after antiretroviral therapy (ART)^45^ to quantify the repertoire changes. RDS decreased with time following ART initiation (ρ = −0.323, *P value=* 6.9 × 10^-6^; Fig. 5I). Consistently, paired comparison of the same patients before and after treatment showed significantly lower RDS after ART (Fig. 5J). These results suggest that immune recovery following ART is accompanied by a gradual rehabilitation of repertoire architecture. The treatment-associated decrease in RDS was observed across multiple V genes, with TRBV27 and TRBV7-3 showing the strongest changes (Fig. 5K). We therefore visualized these two V genes in patients with the most sampled timepoints (n=10). As reported in the original study, Patient 1007 showed a subsequent increase in viral load accompanied by a decline in CD4⁺ T cell counts two years after ART, whereas patient 1024 maintained low viral load and showed recovery of CD4⁺ T cell counts^45^ (Fig. 5L). We therefore examined whether these differences in clinical outcomes were reflected in the TCRmeta space. Indeed, patient 1007 showed limited recovery of reference-associated structures and density peaks over the study period (Fig. 5M). In contrast, patient 1024 gradually recovered distinct local structures and density peaks resembling those in the young healthy reference (Fig. 5M). The agreement between repertoire changes and clinical outcomes illustrates the potential of TCRmeta for patient-level immune monitoring.

### Energy Shift is robust to technical and genetic variation while remaining sensitive to biological repertoire differences

We next sought to evaluate the utilities of TCRmeta in cross-sectional TCR repertoire analysis. Previous approaches, such as the Jaccard index^46^ or Earth Mover’s Distance (EMD) have been introduced to compare repertoire-level distributions^47^. However, they can be heavily influenced by technical factors such as sequencing depth and batch effects, making it challenging to differentiate biological signals from technical variation^5, 9^. To address this issue, we developed a new score, namely “Energy Shift” to measure the separation between two repertoires in TCRmeta space. It is calculated as twice the average TCRmeta distance between the two repertoires minus the average pairwise distance within each repertoire (Fig. 6A). A larger Energy Shift indicates greater separation between the two repertoire distributions.

**Fig. 6.**
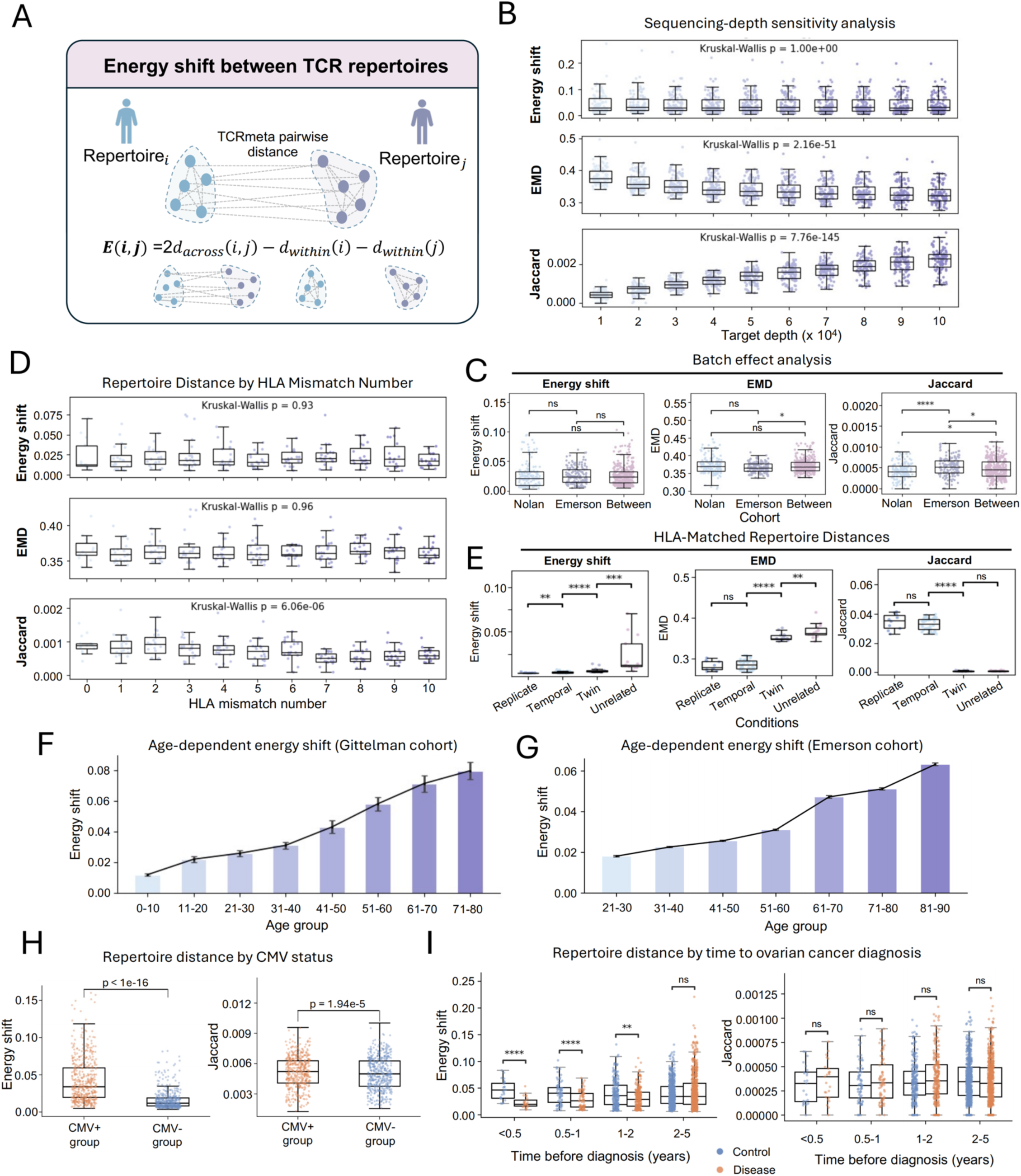
Energy Shift is robust to technical and genetic variation while remaining sensitive to biological repertoire differences. **(A)** Definition of Energy Shift, calculated as twice the mean pairwise TCRmeta distance between two repertoires minus the mean within-repertoire distances of the two repertoires. **(B)** Robustness of Energy Shift, Earth Mover’s Distance (EMD), and Jaccard index to sequencing depth. Repertoires were downsampled from 10,000 to 100,000 TCRs in increments of 10,000. Kruskal-Wallis *P* values across sequencing depths are shown. **(C)** Analysis of batch effects by comparing within-cohort repertoire pairs from the Nolan and Emerson cohorts with between-cohort pairs using Energy Shift, EMD, and Jaccard index. **(D)** Energy Shift, EMD, and Jaccard index between sample pairs with different numbers of human leukocyte antigen (HLA) mismatches. Kruskal-Wallis P values across HLA mismatch groups are shown. **(E)** Energy Shift, EMD, and Jaccard index between HLA-matched technical replicates, temporal samples, twins, and unrelated individuals. Technical replicates represent samples collected from the same individual at the same time point, whereas temporal pairs represent samples collected from the same individual at different time points. **(F, G)** Age-dependent Energy Shift in the **(F)** Gittelman and **(G)** Emerson cohorts. Bars show mean Energy Shift for each age group, and error bars indicate the standard error of the mean. **(H)** Energy Shift and Jaccard index between age-controlled repertoire pairs stratified by CMV status. **(I)** Energy Shift and Jaccard index between age-controlled sample pairs from pre-diagnostic ovarian cancer cases and controls, stratified by time before diagnosis. Statistical significance was assessed using two-sided Mann-Whitney U tests. ns, not significant; *: P value < 0.05; **: P value < 0.01; ***: P value < 0.001; ****: P value < 0.0001.

We first checked the impact of sequencing depth, a common confounder in TCR-seq analysis^48^. We randomly selected 100 repertoire pairs and independently downsampled each repertoire in these pairs to depths ranging from 10,000 to 100,000 TCR counts. The same repertoire pairs were compared at each depth. Energy Shift remained stable across sequencing depths (Kruskal-Wallis *P value=* 1.00), whereas both EMD (*P value=* 2.16 × 10^-^⁵¹) and the Jaccard index (*P value=* 7.76 × 10^-^¹⁴⁵) changed significantly (Fig. 6B).

We next examined batch effects. Differences in experimental procedures can introduce systematic variation between cohorts and bias repertoire comparisons. We compared the Nolan^14^ and Emerson cohorts^7^ while controlling for age and sequencing depth. The two cohorts are considered biologically homogeneous. Eighteen individuals aged 55 years were selected from each cohort, and all repertoires were downsampled to the same depth. We compared repertoire pairs within Nolan (n = 153), within Emerson (n = 153), and between the two cohorts (n = 324). Energy Shift remained similar between within- and between-cohort comparisons, indicating minimal sensitivity to batch effects. In contrast, EMD showed a modest increase in between-cohort distance, whereas Jaccard similarity decreased more significantly between cohorts (Fig. 6C).

The last factor we checked is genetic background. Specifically, differences in HLA typing may contribute to repertoire differences between individuals^49^. We grouped repertoire pairs by the number of HLA mismatches in this analysis. In the Emerson cohort, Energy Shift remained stable across increasing numbers of HLA mismatches (Kruskal-Wallis *P value=* 0.93), as did EMD (*P value=* 0.96), whereas the Jaccard index decreased significantly with increasing HLA mismatch (*P value=* 6.06× 10^-6^; Fig. 6D). The same pattern was reproduced in an independent cohort^50^ (Extended Data Fig. 8A).

The above results collectively indicate that Energy Shift captures repertoire variations beyond technical and genetic factors, and thus likely reflects the environmental changes. Hence, we next investigated whether Energy Shift retained sensitivity to biological differences between repertoires. We compared four types of HLA-matched sample pairs reflecting progressively greater biological differences: When analyzing samples from the same individual^51^, Energy Shift was significantly lower between technical replicates than between longitudinal samples collected at different time points (Fig. 6E), demonstrating its sensitivity to temporal changes in repertoire organization. We then extended the comparison across different individuals. Twins provided an intermediate comparison because they may have distinct antigen exposures yet with identical genetic backgrounds and similar early-life experiences^52^. Twin pairs showed greater Energy Shift than longitudinal samples from the same individuals but remained substantially more similar than HLA-matched unrelated pairs in Emerson cohort (Fig. 6E), showing that Energy Shift can capture biological similarity across individuals. In comparison, EMD did not distinguish technical replicates from longitudinal samples, and the Jaccard index distinguished neither replicates from longitudinal samples nor twins from unrelated individuals. Notably, technical replicates showed less than 4% exact clonotype overlap (Fig. 6E), indicating that resampling alone can introduce substantial variation in the same repertoire. In contrast, their Energy Shift remained close to zero, showing that the overall repertoire organization in TCRmeta space was largely preserved across independent blood draws.

We further compared the sensitivity of Energy Shift and EMD using signal-to-noise ratio (SNR), calculated as the distance between samples with biological differences divided by the distance between technical replicates. Energy Shift showed a higher SNR than EMD for both longitudinal samples (1.2 vs. 1.0) and samples from different individuals (8.5 vs. 1.4; Extended Data Fig. 7A), indicating better separation of biological variation from sampling noise. Energy Shift was also faster to compute than EMD (5.1 vs. 7.8 ms per call; Extended Data Fig. 7B).

Having established the sensitivity of Energy Shift to biological differences, we next examined whether it provides another angle to study immune repertoire aging. Within each age group, we obtained repertoire pairs between individuals in the same age range and calculated their Energy Shift. Energy Shift increased progressively across age groups in the Gittelman cohort (Fig. 6F), and the same pattern was reproduced in the Emerson cohort (Fig. 6G). The increasing Energy Shift with age supports our previous finding that TCR repertoires become more diverse and individualized during aging. In comparison, sex and race had little effect on Energy Shift, with no significant difference between same- and different-sex or same- and different-race repertoire pairs (Extended Data Fig. 8B-C).

Finally, we tested whether Energy Shift could capture repertoire differences associated with disease. We re-analyzed individuals with CMV infection^7^ by selecting age-matched CMV-positive and CMV-negative individuals to limit the effect of age. Energy Shift was significantly higher among CMV-positive individuals than among CMV-negative individuals (Fig. 6H), indicating greater inter-individual repertoire variation in the CMV-positive population. Although the Jaccard index showed a similar trend, the difference between the two groups was substantially more pronounced with Energy Shift. This is consistent with the increased repertoire dispersal observed in CMV-positive individuals in our previous results. We then asked whether Energy Shift could detect more subtle disease-associated differences before cancer diagnosis. We chose ovarian cancer, as it is often asymptomatic at early stages. Our group has previously collected blood samples up to 5 years before ovarian cancer diagnosis together with age-matched healthy controls^53^. Energy Shift was lower among individuals who later developed ovarian cancer than among controls, and the difference became more pronounced when closer to diagnosis (Fig. 6I). In contrast, the Jaccard index showed no difference between the two groups at any time point. Together, these analyses demonstrate that Energy Shift is robust to major technical variation while retaining sensitivity to biological and disease-associated differences in TCR repertoires.

## Discussion

The extensive diversity and individuality of TCR repertoires have long complicated the identification of shared immune patterns across individuals. In this study, we developed TCRmeta to organize diverse TCRs within a common antigen-aware representation and enable repertoire comparison beyond exact sequence identity. Across multiple cohorts and biological settings, TCRmeta revealed conserved repertoire organization despite substantial sequence-level diversity. The Repertoire Dispersal Score (RDS) quantified the deviation of a repertoire from a healthy reference, whereas Energy Shift measured differences directly between repertoires while remaining robust to sampling and technical variation. Together, TCRmeta provides a framework for quantitative cross-individual and longitudinal TCR repertoire analysis.

The performance of TCRmeta likely benefits from integrating the complementary information during its two-stage training. A key feature of pre-training is the use of context-dependent sequence variation rather than independent AA substitutions. We refer to this strategy as neighborhood-conditioned distribution learning (NCDL), in which a bilateral Markov random field models context-dependent AA distributions within TCR neighborhoods. NCDL provides a biologically informed training target that aligns with the contextual representations of BERT-based models, allowing functional motifs associated with related antigen recognition to be learned. Fine-tuning further incorporates experimentally defined antigen relationships and TCR cross-reactivity to organize the embedding space around antigen specificity. NCDL could also be adapted to other sequence-function problems with strong residue dependencies, such as antibody-antigen recognition, enzyme specificity, and protein-protein interactions.

An important implication of our findings is that sequence-level diversity does not necessarily translate into equivalent differences in repertoire organization. This separation may, in part, reflect redundancy in antigen recognition. Genetic variation and stochastic V(D)J recombination generate largely distinct TCR sequences across individuals, yet some different TCRs can recognize the same or related antigens. Such redundancy may allow different individuals to maintain broad coverage of common antigens using different sets of TCR sequences. Consistent with this rationale, technical replicates shared less than 4% of exact TCR clonotypes, yet their Energy Shift remained close to zero (Fig. 6E). This suggests that an individual’s repertoire has a robust underlying organization that can be consistently captured despite incomplete sampling, such that even a limited random subset of clonotypes may reflect broader repertoire structure.

HLA genotype provides another source of sequence-level variation, as it strongly shapes TCR repertoire composition^49^. Individuals sharing HLA alleles present more similar sets of peptides and are therefore more likely to select some of the same TCR clonotypes, consistent with the increase in Jaccard similarity observed with decreasing HLA mismatch (Fig. 6D; Extended Data Fig. 8A). In contrast, Energy Shift showed little dependence on HLA mismatch, suggesting that different HLA backgrounds may select distinct TCR sequences while preserving similar antigen-related repertoire organization. This property enables TCRmeta to compare repertoires across different HLA backgrounds and supports its generalizability for cross-individual repertoire analysis.

The distinct repertoire changes observed across aging, chronic infection and immune reconstitution further suggest that TCRmeta can capture different aspects of repertoire remodeling across biological conditions. The aging-associated changes observed here extend our previous analysis using Repertoire Functional Units (RFUs)^18^. The RFU study identified 13 reproducible age-associated TCR components across cohorts, which provided evidence for conserved repertoire changes with aging. TCRmeta complements this framework by examining the entire repertoire within a continuous antigen-related space, allowing age-associated changes to be evaluated at the level of overall repertoire organization. This analysis revealed a global increase in repertoire dispersal together with reproducible V gene-specific aging patterns. Since V gene information was not explicitly incorporated into RFU representation, TCRmeta provides an additional level of resolution for characterizing age-associated repertoire remodeling.

Our study has several limitations. First, the repertoire-derived TCR relationships used for pre-training provide an indirect approximation of antigen-related relationships but may also reflect factors unrelated to antigen specificity. Second, paired TCR-antigen data remain sparse and unevenly distributed across antigens and HLA types, which limited the antigen-specific information for model training. Third, TCRmeta was primarily developed and evaluated using TCRβ sequences, whereas antigen recognition is determined by paired αβ TCRs. Incorporating the α chain increases the diversity of TCR representations and may provide additional information for distinguishing TCR specificity. Nevertheless, our paired αβ analysis showed that repertoires from different individuals still largely occupied a shared space after α-chain information was incorporated (Fig. 3C), supporting the conserved repertoire organization observed from TCRβ alone. Finally, the clinical utility of TCRmeta has not been fully investigated in this study, which requires future investigations in larger and more diverse populations.

In conclusion, TCRmeta offers a biologically relevant embedding to investigate TCR repertoires at micro- and macro-scales. It reveals shared repertoire organization across individuals despite extensive sequence-level diversity. This framework provides a foundation for cross-individual and longitudinal TCR repertoire analysis and supports the use of TCR repertoires for personalized immune monitoring.

## Materials and Methods

### Pre-training and construction of TCR-specific neighborhoods

TCRmeta was pre-trained using 4,299 TCR repertoire samples combined from two large cohorts^13, 14^. For each repertoire, TCRs were clustered using GIANA algorithm^10^ based on their CDR3 AA sequences and V gene. Similar TCRs found within the same individual were assumed to recognize the same or related antigens and thus have similar antigen specificity. We therefore treated each within-repertoire TCR cluster as a specificity-related group.

For a given TCR, we identified all clusters containing that TCR across all the samples and pooled constituent TCRs to form its neighborhood. Importantly, clusters were not globally merged based on overlapping TCR membership. Instead, each TCR served as an independent anchor and its neighborhood was calculated. Consequently, TCRs from one of the same clusters could have different neighborhoods after this integration process. To ensure sufficient sequence diversity for distribution modeling, only TCRs with neighborhoods with at least 10 distinct TCRs were retained, resulting in 134,664 TCRs. These distributions were used as soft labels to re-train the pre-trained ESM-2 encoder.

### Modeling TCR-specific neighborhood distributions

For each anchor TCR, we modeled the sequence distribution of its neighborhood to generate positional AA probability distributions. TCRs within each GIANA cluster^10^ shared the same CDR3 length and V gene, allowing AA distributions to be estimated directly at each CDR3 position. We evaluated four approaches (positional, first-order Markov chain, second-order Markov chain, Bilateral Markov random field) that use different levels of sequence context to model the AA distribution at a given position.

The positional model estimated the AA distribution independently at each position *i* as

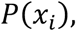

The first-order Markov model conditioned the distribution on the immediately prior residue,

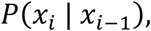

whereas the second-order Markov model considered the two prior residues,

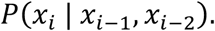

The Bilateral Markov random field model used the adjacent sequence context on both sides of position *i*,

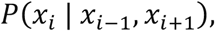

For each anchor TCR, these distributions were estimated empirically from the TCRs in its neighborhood. A specific example of these four modeling approaches was shown in Extended Data Fig. 1B. The frequencies of the AA observed at position *i* among these TCRs were normalized to obtain a conditional probability distribution.

### Soft-label pre-training of the ESM-2 encoder

The neighborhood-derived AA distributions were used as soft labels to re-train the ESM-2^19^ sequence encoder. We used the 35-million-parameter ESM-2 model (esm2_t12_35M_UR50D; embedding dimension, 480), selected after comparison with the 8-million- and 150-million-parameter variants to balance model performance and computational burden. The model input included the CDR3 sequence together with V gene-derived sequence features (CDR1/CDR2/CDR2.5).

Approximately 15% of CDR3 residues were masked for each training sequence. Masking was restricted to internal CDR3 positions, with positions near the two sides sampled at a lower probability than internal positions. Adjacent positions were not masked simultaneously, ensuring that the residues flanking each masked position remained visible to the model for bidirectional context learning.

For each masked position, the model predicted a probability distribution over the 20 standard AAs. Training used a weighted combination of masked-residue cross-entropy and Kullback-Leibler (KL) divergence between the neighborhood-derived target distribution *q* and the model-predicted distribution *p*:

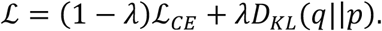

The weight λ was increased linearly from 0 to 1 over the first 100,000 training steps, gradually shifting the training objective from reconstruction of the original AA toward prediction of the neighborhood-derived distribution.

Models were trained using AdamW with a learning rate of 5×10^-5^, batch size of 32, and maximum of 30 epochs. 10% of the data were held out for validation. The learning rate was reduced by a factor of 0.5 when validation loss plateaued, with a minimum learning rate of 10^-7^. Gradient norms were clipped at 1.0, and model checkpoints were selected based on validation loss. After pre-training, the 480-dimensional representation of the [CLS] token was used as the embedding of each TCR.

### Selection and validation of the pre-training strategy

To compare the different pre-training strategies, we first evaluated whether the resulting embeddings placed TCRs from the same GIANA-defined neighborhood closer than TCRs outside the neighborhood. This analysis was performed using an independent Emerson cohort^7^. TCRs overlapping with the pre-training data were excluded, and TCR neighborhoods were constructed using the same GIANA-based procedure as used for pre-training. For each query TCR, positive TCRs were selected from other TCRs within the same GIANA neighborhood. For each positive TCR, control TCRs were selected from outside the neighborhood with the same Hamming distance from the query and mutations at the same CDR3 positions. This design controlled for sequence similarity while testing whether the models could recover GIANA-defined neighborhood relationships.

Euclidean distances were calculated between the query and positive or control TCRs in each embedding space. Neighborhood recovery accuracy was defined as the proportion of positive TCRs whose distance to the query was smaller than the mean distance of the corresponding control TCRs. We additionally calculated a normalized distance margin for each positive-control comparison as

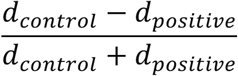

where larger values indicate greater separation of positive and control TCRs relative to the query. The baseline ESM-2 representation was compared with the positional, first-order Markov, second-order Markov chain, and bilateral Markov random field pre-training strategies.

We further evaluated the local antigen organization of each pre-trained embedding using embedding neighborhood consistency. For each model, pairwise Euclidean distances were calculated between TCRs within the same GIANA cluster in the pre-training data, and the 95th percentile of these distances was used as a model-specific distance threshold. The threshold was then applied to an evaluation dataset independent of the pre-training data, assembled from four TCR specificity databases (IEDB^22^, VDJdb^23^, immuneCODE^14^, and McPAS^24^). For each query TCR, embedding neighborhood consistency was defined as the proportion of TCRs within the distance threshold that shared at least one recognized antigen with the query TCR.

### Construction of antigen-related TCR pairs

TCR-antigen binding pairs were collected from IEDB^22^, VDJdb^23^, immuneCODE^14^, and McPAS^24^. TCRs were restricted to those with recognized V genes and CDR3 length of 10-24 AAs. Peptide sequences were required to be 7-19 AAs in length. For VDJdb, only records with a confidence score greater than one were used.

Rather than defining two TCRs as a positive pair based on one shared binder, we evaluated the similarity of their annotated binder pools. For each TCR pair, a 2×2 contingency table was constructed from the number of peptides recognized by both TCRs, by either TCR alone, or by neither TCR (Extended Data Fig. 1A). To reduce the contribution of highly represented antigens, peptides were down-weighted according to their frequency in the training data, as highly dominant peptides may contain a greater proportion of false-positive annotations.

One-sided Fisher’s exact tests were used to identify enrichment or depletion of binder pool. Enrichment (alternative = “greater”) was used to define positive TCR pairs, whereas depletion (alternative = “less”) was used to define negative pairs. The resulting dataset contained 3,007,593 significant positive TCR pairs and 9,790,950 significant negative TCR pairs.

Importantly, neither antigen identity nor antigen sequence was provided as input to the model. Fine-tuning was performed entirely on TCR representations, with antigen annotations serving only as supervision for learning antigen-associated TCR relationships.

### Fine-tuning of TCR representations

The pre-trained TCR representations were further fine-tuned using antigen-informed TCR pairs. We designed the fine-tuning objective to explicitly organize the geometry of the embedding space, such that TCRs with related antigen-binding profiles were positioned close together while unrelated TCRs were separated.

Each 480-dimensional TCR representation was passed through a shared three-block residual multilayer perceptron (ResMLP) with a hidden dimension of 768 and dropout of 0.3, followed by projection to a 64-dimensional representation and *L_2_* normalization. For a pair of TCRs with projected representations *z_i_* and *z_j_*, their distance was defined as the scaled Euclidean distance

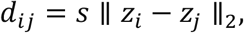

where *s* is a learnable scaling parameter. Pairwise distance was converted to a classification logit *P_i_*_j_ using a learnable distance threshold and scaling factor *α*,

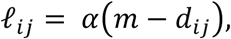

such that antigen-associated TCR pairs were encouraged to have shorter distances.

To provide a continuous measure of similarity between the antigen-binding profiles of two TCRs, we defined antigen distances. For each TCR, all annotated peptide binders were encoded using a baseline ESM-2 model, and the centroid of their peptide embeddings was calculated. *P_i_* denotes the set of peptide binders for TCR *i*, its antigen center was defined as

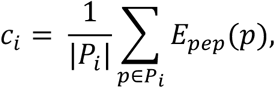

where *E_pep_*(*p*) is the ESM-2 representation of peptide *p*. The antigen distance between two TCRs was then calculated as the Euclidean distance between their peptide centers

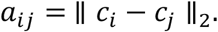

The final fine-tuning objective combined weighted binary cross-entropy, ranking, top-k ranked purity, and contrastive losses:

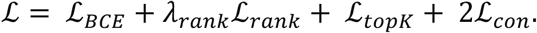

For *N* TCR pairs, the weighted binary cross-entropy loss was defined as

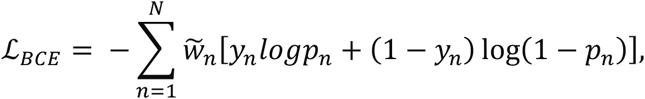

where *y_n_* is the antigen-informed pair label and *w*A*_n_* is the normalized pair weight. Negative pairs were assigned an additional twofold weight relative to positive pairs.

To preserve structural antigen-related relationships in the embedding space, the ranking loss compared TCR pairs according to their antigen-derived distances. For pairs *p* and *q* with antigen distances *a_p_* > *a_q_*,

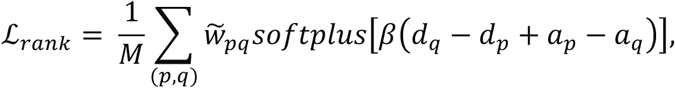

where *β*=10 and *W~_pq_* incorporates the difference in antigen distance and pair-specific weights. The ranking loss therefore encouraged TCR pairs with more similar antigen-binding profiles to remain closer in the learned representation space. Ranking was performed both within individual batches and against historical pairs stored in a memory buffer to provide cross-batch comparisons. The ranking contribution was introduced after the first training epoch and increased linearly to its final weight by the third epoch.

To improve the purity of local TCR neighborhoods, an additional loss was applied to the K pairs with the smallest embedding distances:

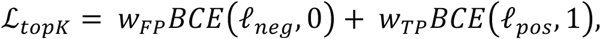

With

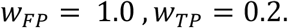

Thus, false-positive negative pairs appearing among the closest TCRs were penalized fivefold more strongly than the reward assigned to correctly positioned positive pairs. This asymmetry was introduced to prioritize specificity and reduce false-positive relationships within local TCR neighborhoods.

Finally, a contrastive loss directly constrained pairwise embedding distances,

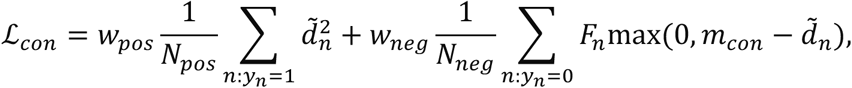

where *d~_n_ = d_n_/d_max_* is the normalized TCR distance, *m_con_* is the negative-pair margin, and *F_n_* provides additional weighting according to the proximity of a negative pair to the margin. Together, these objectives encouraged antigen-related TCRs to form locally coherent neighborhoods while separating antigen-unrelated TCRs and reducing false-positive relationships among nearby TCRs.

Models were trained using AdamW with a learning rate of 1×10^-5^, weight decay of 1×10^-4^, a batch size of 256, and up to 50 epochs. Model parameters were maintained using an exponential moving average with a decay of 0.999. The learning rate was reduced by a factor of 0.5 when validation loss plateaued, with a minimum learning rate of 10^-7^. Gradient norms were clipped at 5.0.

### Validation using TCR deep mutational scanning datasets

We evaluated pre-trained TCRmeta using five published TCR deep mutational scanning (DMS) studies comprising seven TCRs: A3, DMF4, 868-Z11, NLV3, A6c134, RD1, and S3-4 (Fig. 2B; Extended Data Fig. 2A). These datasets provided 12 experimental readouts in total, including both binding and activation measurements for A3 and DMF4^27^ and binding measurements against four different antigens for S3-4^28^. For two studies in which numerical DMS measurements were not directly available^28, 29^, fitness scores were inferred from the published figures. For each mutant TCR, the fitness score was defined as the absolute change in enrichment relative to the wild-type (WT) TCR. TCR distance was calculated as the Euclidean distance between mutant and WT embeddings in each model. Spearman correlation was used to assess the association between TCR distance and the absolute fitness score at each position. To examine whether performance varied across sequence features, correlations were further summarized by the CDR3 position.

### Benchmark on independent TCR-antigen specificity database

TCRmeta was benchmarked using four independent TCR antigen-specificity datasets: a held-out subset from same source as the training dataset, TetTCR-Seq^30^, TetTCR-SeqHD^31^, and TChard^32^. To ensure no similar TCRs in the held-out subset (data leakage), training and test TCRs were jointly clustered using GIANA, and test TCRs within the same cluster with any training TCR were excluded. For the three external datasets, exact CDR3 and V gene overlaps with the training data were removed.

Positive and negative TCR-TCR pairs were constructed independently for each dataset using the same Fisher’s exact test procedure used for fine-tuning. This resulted in 14,720, 6,300, 5,812, and 68,914 TCR pairs for the held-out, TetTCR-Seq, TetTCR-SeqHD, and TChard datasets, respectively. To facilitate comparison, positive and negative pairs were balanced by retaining the most significant pairs from the larger class.

TCRmeta was compared with TCR-VALID^15^, CDR3vec^16^, SCEPTR^17^, TCR-BERT^11^, GIANA^10^, RFU^18^, ESM-2^19^, and TCRdist^6^ using the same TCR pairs. Performance was evaluated by top-K% purity. TCR pairs were ranked by increasing distance in each representation space, and purity was calculated as the fraction of positive pairs among the top closest pairs,

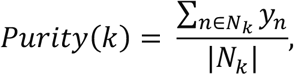

where *N_k_* denotes the top k% of pairs with the smallest distances and *y_pos_* = 1, *y_ne_*_A_ = 0. This metric evaluates the enrichment of antigen-related TCRs among the closest pairs and how well this specificity is maintained as the distance cutoff progressively expanded.

To determine whether the enrichment observed with TCRmeta was greater than expected by chance, a permutation test was performed independently for each dataset. TCRmeta distances were kept fixed, while positive and negative pair labels were randomly permuted while preserving the original number of pairs in each class. For each permutation, the mean top 1-30% purity was calculated. A total of 5,000 permutations were performed, and a one-sided empirical *P* value was calculated by comparing the observed mean purity with the permutation distribution using a +1 correction.

Bootstrap resampling was further used to quantify differences between TCRmeta and each comparison method. For each dataset, TCR pairs were sampled with replacement to generate 5,000 bootstrap replicates. Within each replicate, the mean top 1-30% purity was calculated for TCRmeta and each comparison method, and the difference was defined as TCRmeta purity minus comparison-method purity. The median bootstrap difference was reported together with the 2.5th and 97.5th percentiles. Positive values indicate higher antigen-related pair purity for TCRmeta

### Embedding-space visualization of TCR repertoires across cohorts and disease states

To visualize TCR repertoire organization across cohorts and biological conditions, we compared TCRmeta embeddings from healthy individuals of different age groups (0-1^33^, 25-30^14^, and 86-89 years^14^), individuals with type 1 diabetes (T1D)^33^, lung cancer^34^, or RAG mutations^35^, and thymocytes^36^. Most groups were obtained from different cohorts. We also generated 10,000 random TCR-like sequences as a negative control. Each sequence contained 8-13 randomly sampled AAs flanked by cysteine (C) and phenylalanine (F).

The RAG mutation group had the fewest samples and TCR clonotypes and was therefore used as the reference for downsampling. Eight samples were included from each biological group. The number of TCR clonotypes was also matched across groups, with 47,689 clonotypes per group across all TRBV genes. For the three representative V genes, each group contained 8,739 TRBV28, 3,644 TRBV27, and 2,962 TRBV6-5 clonotypes. This matching ensured that differences in the visualized distributions were not driven by differences in sample or clonotype numbers. TCRmeta embeddings were projected into two dimensions using t-SNE (perplexity = 30, PCA initialization, and automatic learning rate). TCRs were colored by CDR3 length to visualize sequence-length organization within the embedding space.

### Evaluation of shared embedding structure using α and paired αβ TCRs

Paired TCR α- and β-chain sequences were obtained from the Observed TCR Space (OTS) database^37^, including healthy, COVID-19, cancer, and autoimmune samples. The same set of paired clonotypes was analyzed using α-chain, β-chain, or concatenated αβ embeddings. TCRs were grouped by TRAV-TRBV combination, retaining groups with at least three samples and five clonotypes per sample. Clonotype numbers were matched across samples within each group.

For each TRAV-TRBV combination, mean Euclidean distances were calculated between TCRs from the same individual (within-individual) and from different individuals (cross-individual). The cross/within distance ratio was calculated to compare individual-level separation in the embedding space. The same analysis was performed using α-only, β-only, and paired αβ representations.

### TCR repertoire downsampling

To control for differences in sequencing depth, TCR repertoires were downsampled to a fixed target depth using multinomial sampling. For each repertoire, sampling was based on template counts, so that more abundant clonotypes were more likely to be sampled. Repertoires with total counts below the target depth were kept without downsampling. Unless otherwise specified, all downsampling in this study followed this procedure, with a target depth of 10,000 TCR counts.

### Construction of the healthy repertoire reference map

A healthy repertoire reference was constructed using 500 individuals aged 0-30 years from the Gittelman cohort^13^, including 99 individuals aged 0-10, 169 aged 11-20, and 232 aged 21-30 years. Samples were restricted to the 2-month (first time point) visit and to individuals who were COVID-negative, unvaccinated, and asymptomatic. Each repertoire was downsampled to 10,000 TCR counts to balance the contribution of individual samples. TRBV genes with a pooled frequency of at least 1% were retained and modeled separately (Extended Data Fig. 5A).

For each TRBV gene, TCRmeta embeddings from the reference samples were pooled and reduced by principal component analysis (PCA) before density estimation. We evaluated different numbers of PCA dimensions by comparing the retained explained variance, agreement with density estimates from the full 64-dimensional embedding space, and the stability of the association between RDS and age (Extended Data Fig. 5B). The age association increased with the number of dimensions and largely stabilized around 30 dimensions. At 30 dimensions, density estimates showed high agreement with those obtained from the full embedding space (Spearman ρ = 0.98) while retaining 91.1% of the explained variance. We therefore used 30 PCA dimensions for subsequent analyses. A Gaussian kernel density estimator (KDE) was then fitted to the PCA-transformed embeddings, weighted by clonotype abundance. The KDE bandwidth was set as *n*^−1/(*d*+4)^, where *n* is the number of clonotypes and *d* is the number of PCA dimensions. The resulting density models were used as the young healthy reference for evaluating new TCR repertoires.

### Definition of repertoire dispersal score (RDS)

Before RDS calculation, each repertoire was downsampled to 10,000 TCR counts by multinomial sampling according to clonotype abundance. Thus, each clonotype retained a downsampled count representing its abundance in the standardized repertoire.

For each sample, TCRmeta embeddings were mapped to the corresponding TRBV-specific healthy reference density model, and each clonotype was assigned a KDE log-density value. For each TRBV gene, low-density (*LD*_15_) and high-density (*HD*_85_) regions were defined using the 15th and 85th percentiles of the healthy reference density distribution, respectively. The LD and HD fractions were calculated as the fractions of downsampled TCR counts assigned to these regions:

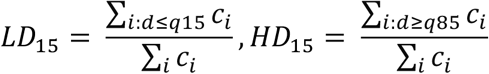

where *d_i_*is the KDE log-density assigned to clonotype *i*, *c_i_*is its downsampled count, and *q*_15_and *q*_85_are the 15th and 85th percentile density thresholds of the corresponding healthy reference model. The TRBV-specific repertoire dispersal score (RDS) was then defined as

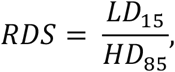

A sample-level RDS was obtained by averaging the TRBV-specific scores across V genes, weighted by the number of unique clonotypes represented within each V gene. Higher RDS values indicate greater repertoire dispersal from the healthy reference.

### Visualization of TCRmeta embeddings

TCRmeta embeddings were visualized in two dimensions using t-distributed stochastic neighbor embedding (t-SNE) or Uniform Manifold Approximation and Projection (UMAP). For visualization of overall repertoire organization (Fig. 3A), the 64-dimensional TCRmeta embeddings were directly projected into two dimensions using t-SNE. For healthy reference-based analyses, UMAP was used to establish a common two-dimensional reference map, allowing additional repertoires to be projected into the same space. Sample embeddings were transformed using the scaler fitted to the corresponding TRBV-specific healthy reference and projected using the fitted UMAP reducer.

### MAIT T cells analysis

Human MAIT TCR sequences were collected from five published datasets^39, 40, 41, 42, 43^. MAIT TCRs were embedded using TCRmeta and analyzed separately by V gene. For each V gene, MAIT TCRs and TCRs from different age groups were projected onto the same UMAP space using the healthy reference. A density map was generated for the MAIT TCRs using KDE. The MAIT region was defined by areas with at least 5% of the peak MAIT density. For each age group, a separate density map was generated using TCR template counts as weights, and the top 5% highest-density region was defined as the high-density (HD) region. The overlap between the HD region and the MAIT region was calculated as the fraction of the HD region overlapping the MAIT region. The association between MAIT overlap and age was evaluated using Spearman correlation.

### Analysis of repertoire dispersal across immune states and interventions

Repertoire dispersal was evaluated across independent aging, CMV, bone marrow transplantation and HIV cohorts using the healthy reference and RDS defined above. For the Gittelman aging cohort, samples were restricted to the 2-month visit and individuals who were COVID-negative, unvaccinated, and asymptomatic. 100 samples were then selected randomly from each age group. While most samples in the 21-30-year age group were used to construct the healthy reference, 10 samples were held out for evaluation. For the Emerson cohort, up to 30 individuals were selected from each age group, prioritizing individuals closest to the midpoint of each age interval. No additional sample filtering was performed for the other datasets.

For each dataset, associations were first evaluated separately for each TRBV gene. The two V genes with the largest absolute Spearman correlation coefficients were selected for visualization.

### Definition of energy shift

Energy Shift was used to quantify the difference between two TCR repertoires in TCRmeta embedding space. Repertoires were first downsampled to 10,000 TCR counts and analyzed separately by TRBV gene. Only V genes containing at least 100 clonotypes in both samples were included. For each V gene, pairwise Euclidean distances were calculated between TCRs from the two repertoires and within each repertoire, using downsampled template counts as weights. Energy Shift was defined as

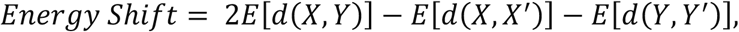

where *X* and *Y* represent the two repertoires. Higher Energy Shift values indicate greater separation between the two repertoire distributions in TCRmeta embedding space.

### Calculation of Jaccard index and Earth Mover’s Distance (EMD)

The Jaccard index and Earth Mover’s Distance (EMD) were used as additional measures for pairwise repertoire comparison. Repertoires were first downsampled to 10,000 TCR counts and analyzed separately by TRBV gene. Only V genes with at least 100 clonotypes in both repertoires were included.

For each V gene, the Jaccard index was calculated based on exact clonotype overlap between two repertoires:

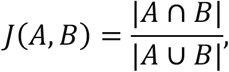

where and represent the sets of unique TCR clonotypes in the two repertoires. Larger Jaccard index indicate greater similarity between the two repertoires.

EMD was calculated using TCRmeta embeddings. Pairwise Euclidean distances between TCRs from the two repertoires were used as the transport cost, with downsampled template counts used as weights. The distance matrix was normalized by its maximum value, and EMD was calculated as the minimum cost required to transform one weighted TCR distribution into the other. Lower EMD values indicate greater similarity between the two repertoires in TCRmeta embedding space.

### Selection of repertoire sampling strategy

To determine an appropriate strategy for controlling repertoire size in pairwise repertoire comparisons, we compared count-based multinomial downsampling with random sampling of unique clonotypes. Multinomial downsampling preserves the original abundance distribution by sampling clonotypes according to their template counts, whereas random clonotype sampling gives each unique clonotype an equal probability of being selected regardless of abundance.

Thirty repertoires with high clone counts were selected for this analysis. For each repertoire, independently resampled repertoires were generated using either multinomial downsampling (weighted) or random sampling of unique clonotypes (uniform). We also evaluated whether sampled clonotypes retained their original abundance (expanded) or were represented once regardless of their original counts (unexpanded). This resulted in four sampling strategies: weighted, uniform, expanded only, and unexpanded only.

We first quantified the variation introduced by resampling. For each original repertoire, two independently resampled repertoires were generated using each pair of sampling strategies, and Energy Shift between the two resampled repertoires was calculated as within-sample Energy (Extended Data Fig. 9A). Lower within-sample Energy indicates less variation introduced by sampling.

We then examined how sampling-induced variation compared with differences between independent repertoires. All pairwise combinations of the 30 repertoires were analyzed, resulting in 435 repertoire pairs. For each pair, the two repertoires were independently processed using the corresponding sampling strategies, and their Energy Shift was calculated as cross-sample Energy (Extended Data Fig. 9B). The signal-to-noise ratio (SNR) was calculated by dividing the mean cross-sample Energy by the mean within-sample Energy for each combination of sampling strategies (Extended Data Fig. 9C). A higher SNR indicates greater separation between independent repertoires relative to the variation introduced by resampling.

Although the expanded-only approach achieved the highest SNR (12.08), its application substantially reduced the available repertoire size in samples with lower sequencing depth. Weighted multinomial downsampling achieved the second-highest SNR (4.34) while retaining more TCRs across repertoires with different sequencing depths. We therefore used weighted multinomial downsampling for subsequent analyses.

### Robustness of Energy Shift to sampling, technical, and genetic variation

To evaluate sensitivity to sequencing depth, we first identified 695 repertoires with more than 100,000 TCR counts in Nolan cohort^14^. All possible pairs among these eligible repertoires were generated, and 100 repertoire pairs were randomly selected without replacement (random seed = 42). The same 100 repertoire pairs were used at all sequencing depths, allowing changes in each metric to be evaluated without differences in sample composition across depth groups. For each pair, both repertoires were independently downsampled to 10,000, 20,000, 30,000, and up to 100,000 TCR counts. Energy Shift, EMD, and the Jaccard index were calculated for each pair at each depth.

To evaluate batch effects, we compared repertoires from the Nolan^14^ and Emerson^7^ cohorts. To minimize biological variation between cohorts, the analysis was restricted to individuals aged 55 years in both datasets. Eighteen repertoires from each cohort were included in the final analysis, and all repertoires were downsampled to 10,000 TCR counts. Three groups of repertoire pairs were then constructed: all pairwise comparisons among the 18 Nolan repertoires (153 Nolan-Nolan pairs), all pairwise comparisons among the 18 Emerson repertoires (153 Emerson-Emerson pairs), and all cross-cohort comparisons between the 18 Nolan and 18 Emerson repertoires (324 Nolan-Emerson pairs). Energy Shift, EMD, and the Jaccard index were calculated for every pair and compared across the three groups. This design allowed cohort-associated differences to be evaluated while holding age and sequencing depth constant.

The effect of HLA background was evaluated in two independent cohorts^7, 50^. Repertoire pairs were grouped according to the number of HLA mismatches between the two individuals. To reduce potential confounding by age, pairs were selected to maintain similar mean ages across HLA mismatch groups. In the Emerson cohort, the final analysis included 213 repertoire pairs from 116 individuals, consisting of 13 pairs with zero HLA mismatches and 20 pairs for each mismatch level from 1 to 10. In the independent validation cohort, 20 repertoire pairs were included for each HLA mismatch level from 2 to 10, resulting in 180 pairs. The same three repertoire distance metrics were calculated for each pair and compared across HLA mismatch groups.

### Sensitivity of Energy Shift to biological repertoire differences

The sensitivity of Energy Shift to biological repertoire differences was evaluated across multiple biological settings. To examine subtle repertoire differences while minimizing the influence of HLA variation, HLA-matched repertoire pairs were classified as technical replicates^51^, longitudinal samples from the same individual^51^, twins^52^, or unrelated individuals^7^. The analysis included 10 technical replicate pairs, 12 longitudinal pairs, and 15 twin pairs and 13 pairs of HLA-matched unrelated individuals. Energy Shift, EMD, and Jaccard index were calculated and compared across the four groups.

Age-associated repertoire differences were evaluated independently in the Gittelman and Emerson cohorts. In the Gittelman cohort, up to 100 individuals were included per 10-year age group, while 30 individuals per age group were selected in the Emerson cohort. Pairwise Energy Shift was calculated within each age group. The analysis covered age groups from 0-10 to 71-80 years in the Gittelman cohort and from 21-30 to 81-90 years in the Emerson cohort.

CMV-associated repertoire differences were evaluated in the Emerson cohort. Because age can affect repertoire differences, CMV-positive and CMV-negative individuals were selected from their overlapping age range of 5-70 years to reduce age-related bias. A total of 100 CMV-positive and 100 CMV-negative individuals were included, with median ages of 42 and 40 years, respectively. Pairwise Energy Shift was calculated separately within the CMV-positive and CMV-negative groups, resulting in 4,950 repertoire pairs per group. Jaccard index was calculated using the same comparison framework.

Ovarian cancer-associated repertoire differences were evaluated using a matched case-control cohort. Samples were grouped according to time before ovarian cancer diagnosis (<0.5, 0.5-1, 1-2, and 2-5 years). Controls were matched to cases using the cohort matching information, and matched pairs with a difference in blood-draw age greater than 2 years were excluded to further control for age. Within each time interval, pairwise Energy Shift was calculated separately for ovarian cancer cases and controls, with Jaccard index calculated for comparison.

### Statistical analysis

Statistical analyses were performed using Python (version 3.10.4). Comparisons between two independent groups were performed using two-sided Mann-Whitney U tests, and comparisons across three or more independent groups were performed using Kruskal-Wallis tests. Associations between continuous or ordinal variables were assessed using two-sided Spearman rank correlation. For analyses performed across multiple TRBV genes, *P* values were corrected for multiple testing using the Benjamini-Hochberg false discovery rate (FDR) method.

For construction of positive and negative TCR pairs used for model fine-tuning, enrichment and depletion were assessed using Fisher’s exact test. A one-sided greater alternative was used to identify positive pairs, whereas a one-sided less alternative was used to identify negative pairs. Box plots in all the figures show the median and interquartile range (IQR), with whiskers extending to 1.5 × IQR.

All statistical tests were two-sided unless a directional alternative was explicitly specified. P < 0.05 was considered statistically significant unless otherwise indicated. Statistical analyses and data processing were performed using NumPy (version 1.26.4), pandas (version 2.2.3), SciPy (version 1.15.3), and scikit-learn (version 1.6.1). Data visualization was performed using Matplotlib (version 3.9.4) and seaborn (version 0.13.2).

## Acknowledgments

Funding

- National Cancer Institute grant R01 CA258524-01A1
- National Cancer Institute grant R01 CA245318-01A1

## Author contributions

Conceptualization: MP, BL

Methodology: MP

Software: MP, YW

Validation: MP

Formal analysis: MP

Investigation: MP, JH

Resources: MP, BL

Data curation: MP

Writing - original draft: MP

Writing - review & editing: MP, BL, YT, YW, JH, HH, JF

Visualization: MP

Supervision: BL

Project administration: BL, MP

Funding acquisition: BL

## Data and materials availability

The TCR datasets used for model training were obtained from the Immune Epitope Database (IEDB; https://www.iedb.org/), VDJdb (https://vdjdb.cdr3.net/), the immuneCODE COVID-19 dataset available through immuneACCESS (https://clients.adaptivebiotech.com/pub/covid-2020), and McPAS-TCR (https://friedmanlab.weizmann.ac.il/McPAS-TCR/). Deep mutational scanning and mutational TCR datasets are available through the respective publications and associated source data (DOIs: 10.1016/j.immuni.2024.11.009, 10.3389/fimmu.2024.1345368, 10.1016/j.immuni.2022.09.004, 10.1021/acs.biochem.0c00689, and 10.1016/j.str.2016.04.011). External datasets used to evaluate antigen-related TCR relationships included TetTCR-Seq (DOI: 10.1038/nbt.4282), TetTCR-SeqHD (DOI: 10.1038/s41590-021-01073-2), and TChard (Zenodo: 10.5281/zenodo.6962043). Population-level TCR repertoire datasets spanning diverse biological and clinical conditions were obtained primarily from the Adaptive Biotechnologies immuneACCESS database, including datasets of SARS-CoV-2 infection, CMV exposure, childhood development, RAG deficiency, lung cancer, HIV infection and antiretroviral therapy, human thymic T cells, and monozygotic twins (DOIs: 10.1172/jci.insight.151849, 10.21417/B7VC7S, 10.21417/ADPT2020COVID, 10.1172/jci.insight.161885, 10.1016/j.molimm.2020.09.003, 10.1016/j.jaci.2017.08.001, 10.1158/2326-6066.CIR-24-1109, 10.21417/VN2025CIR1, 10.3389/fimmu.2022.879190, 10.7554/eLife.63502, and 10.3389/fimmu.2021.599133). Paired TCRαβ repertoire data were obtained from the Observed T Cell Receptor Space (OTS) database (DOI: 10.1016/j.celrep.2024.114704). The ovarian cancer TCR-seq dataset is available through Zenodo (DOI: 10.5281/zenodo.11204147). Additional single-cell TCR datasets used for the MAIT cell analysis were obtained from previously published studies (DOIs: 10.1073/pnas.1902649116, 10.1038/s41586-018-0694-x, 10.1016/j.cell.2017.05.035, 10.1038/s41591-018-0045-3, and 10.1172/jci.insight.147474). Code for using TCRmeta is publicly available on GitHub (https://github.com/Mia-yao/TCRmeta) and PyPI (https://pypi.org/project/TCRmeta/).

