## Supplemental figures for "Antigen-aware TCR embeddings reveal conserved immune patterns for personalized immune monitoring"

**Extended Data Figures**

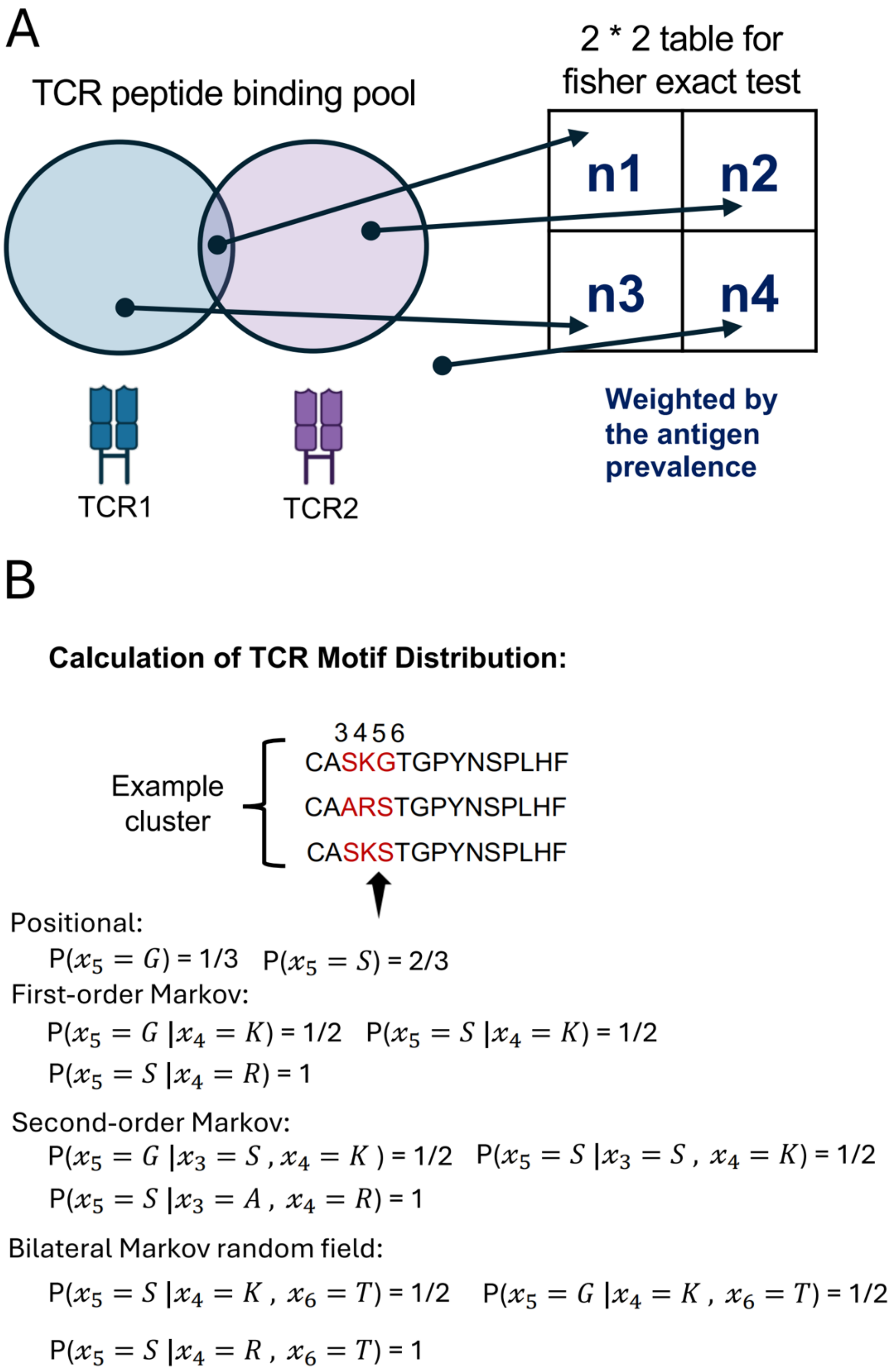


**Extended Data Fig. 1. Additional details of TCRmeta pre-training, benchmarking, and fine-tuning.**

**(A)** Construction of the 2 × 2 contingency table used in Fisher's exact test for each TCR pair during fine-tuning. Counts represent peptides recognized by both TCRs, by either TCR alone, or by neither TCR. Peptides were down-weighted according to their prevalence in the training data to reduce the contribution of highly represented antigens (see Methods). One-sided Fisher's exact tests were used to identify enrichment or depletion of shared binders for defining positive and negative TCR pairs, respectively. **(B)** Example calculation of the TCR neighborhood AA distributions used for pre-training. For a given position within a TCR neighborhood, AA probabilities were calculated using positional, first-order Markov, second-order Markov chain, or bilateral Markov random field based on the AA observed at the corresponding positions.

A

| TCR | CDR3β | Length | V gene | Antigen | Class | Assay | Cohort |
|---|---|---|---|---|---|---|---|
| A3 | CASSPNMADEQYF | 13 | TRBV5-1 | MAGE-A3 | Tumor | Activation, binding | Vazquez-Lombardi |
| DMF4 | CAISEVGVGQPQHF | 14 | TRBV10-3 | MART-1 | Tumor | Activation, binding | Vazquez-Lombardi |
| 868-Z11 | CASSDTVSYEQYF | 13 | TRBV5-6 | HIV SL9 | Viral | Binding | Rosenberg |
| NLV3 | CASSPTGGSPSPLHF | 15 | TRBV27 | HCMV NLV | Viral | Binding | Abdelfattah |
| A6-c134 | CASRPGLMSAQPEQYF | 16 | TRBV6-5 | Tax | Viral | Binding | Harris |
| RD1 | CASRPGWMAGGVEQYF | 16 | TRBV6-5 | MART-1 | Tumor | Binding | Harris |
| S3–4 | CASRPGAAGGRPELYF | 16 | TRBV6-5 | Tax / MART-1 / WT1 / Survivin | Mixed | Binding | Singh |

B

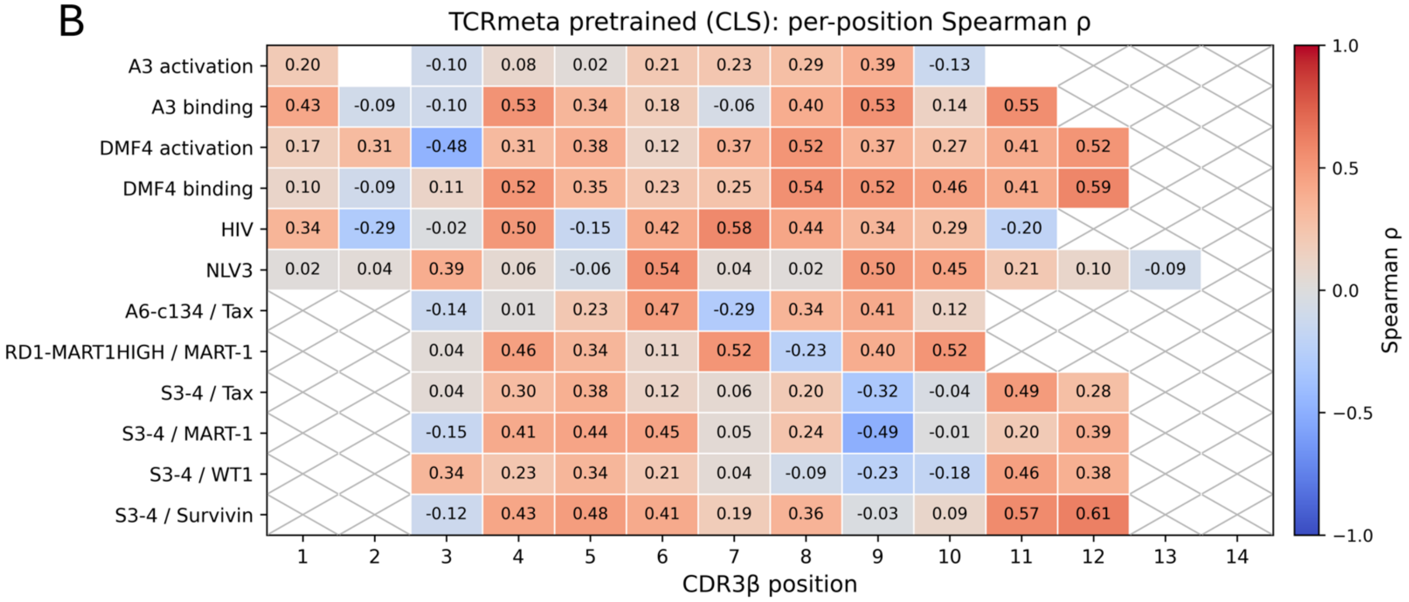


**Extended Data Fig. 2. Deep mutational scanning datasets and position-specific performance of pre-trained TCRmeta.**

**(A)** Summary of the seven TCRs included in the deep mutational scanning (DMS) benchmark, including CDR3β sequence, CDR3β length, V gene, antigen, antigen class, experimental assay, and source study. The benchmark comprises 12 experimental readouts of TCR-pMHC binding or T cell activation from five published DMS studies. **(B)** Position-specific Spearman correlations between mutant-WT distances in the pre-trained TCRmeta embedding space and experimental fitness scores for each DMS readout. Each value represents the Spearman correlation across amino-acid substitutions at the indicated CDR3β position. Crossed cells indicate positions without experimental fitness scores, whereas empty cells indicate positions where all evaluated mutants had the same fitness score and a correlation therefore could not be calculated.

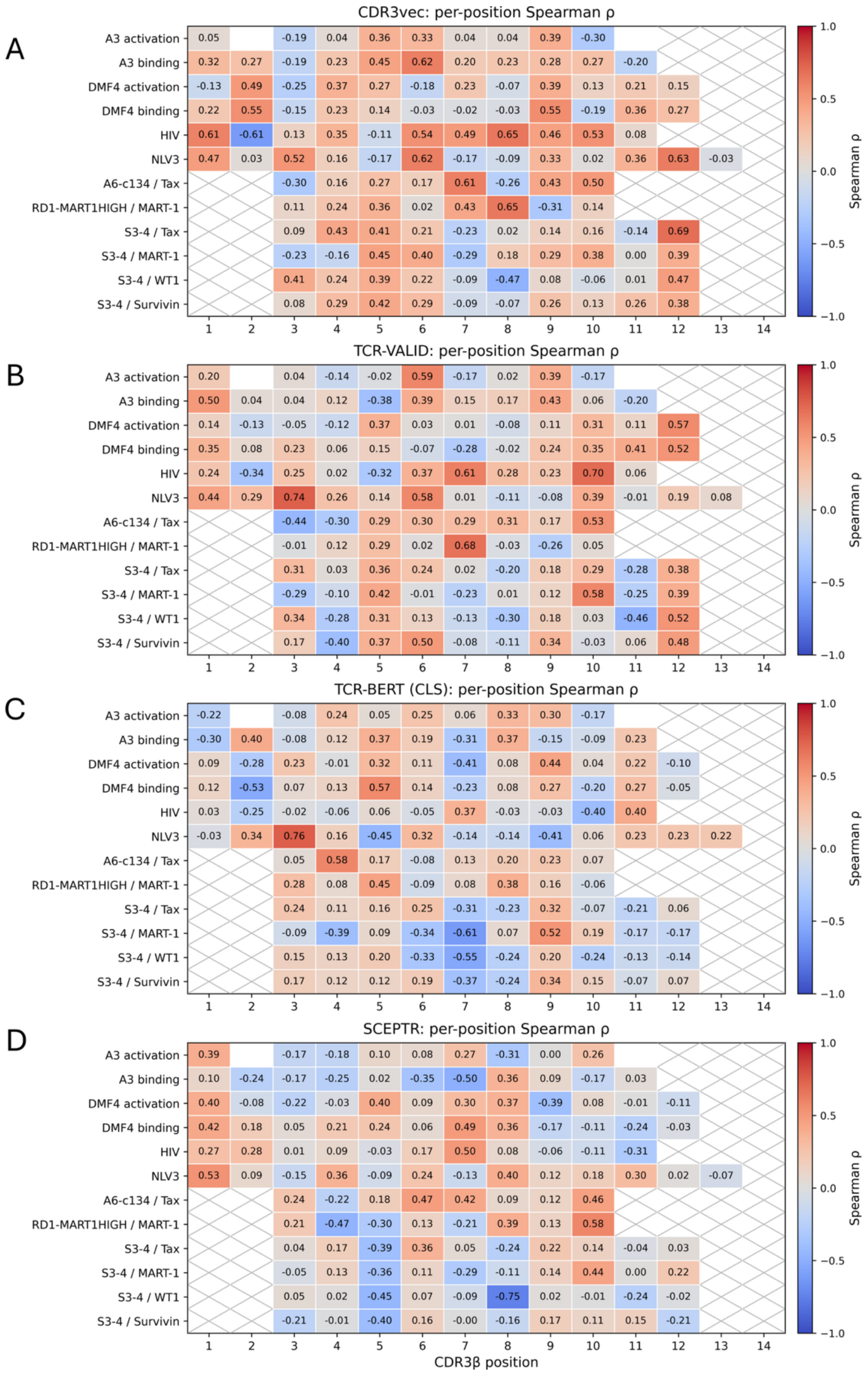


**Extended Data Fig. 3. Position-specific performance of comparison TCR representation models in DMS datasets.**

**(A-D)** Position-specific Spearman correlations between mutant-WT embedding distances and experimental fitness scores for **(A)** CDR3vec, **(B)** TCR-VALID, **(C)** TCR-BERT, and **(D)** SCEPTR across the 12 deep mutational scanning (DMS) readouts. Each value represents the Spearman correlation across amino-acid substitutions at the indicated CDR3β position. Crossed cells indicate positions without experimental fitness scores, whereas empty cells indicate positions where all evaluated mutants had the same fitness score and a correlation therefore could not be calculated.

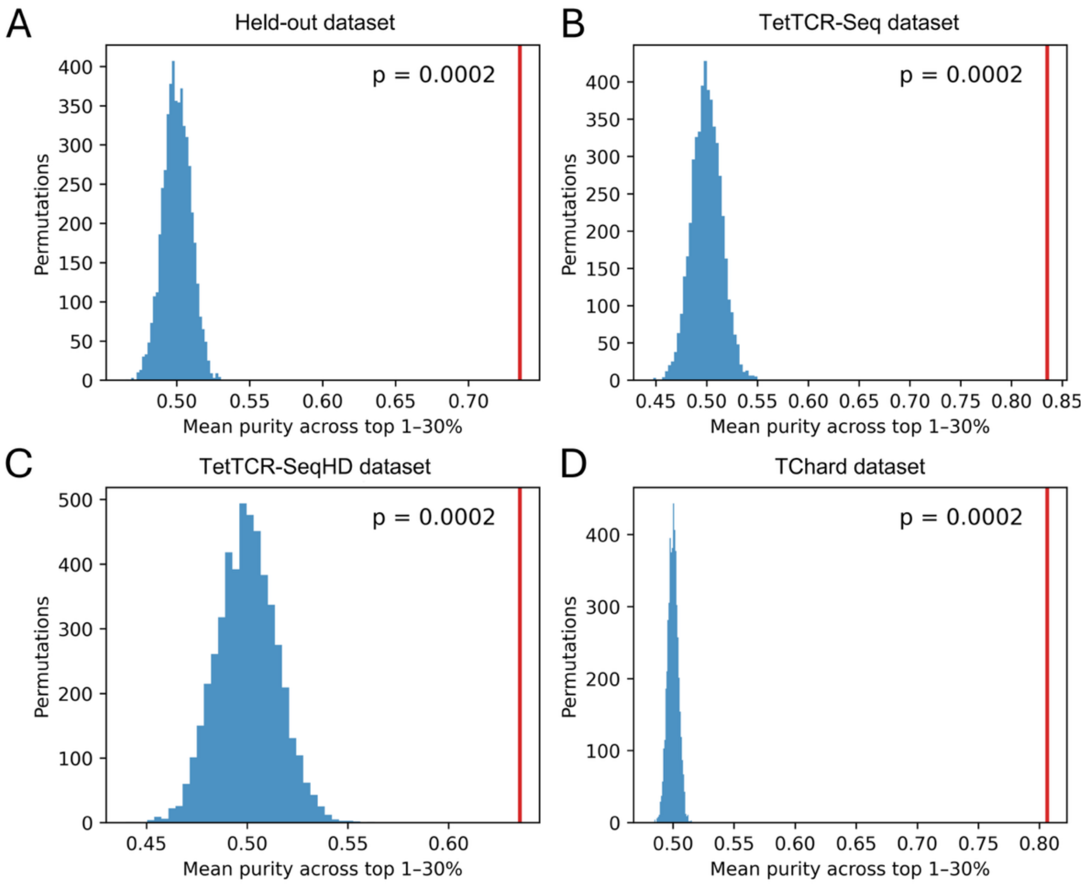


**Extended Data Fig. 4. Permutation analysis of TCRmeta pairwise antigen-related TCR identification.**

**(A-D)** Permutation analysis of TCRmeta performance in the **(A)** held-out**, (B)** TetTCR-Seq, **(C)** TetTCR-SeqHD, and **(D)** TChard datasets. TCR pairs were ranked by TCRmeta embedding distance, and performance was quantified as the mean purity of antigen-related TCR pairs across the top 1-30% nearest pairs. Pair labels were randomly permuted 5,000 times while preserving the numbers of antigen-related and unrelated pairs to generate the null distribution. Red lines indicate the observed TCRmeta values. Empirical one-sided P values were calculated from 5,000 permutations.

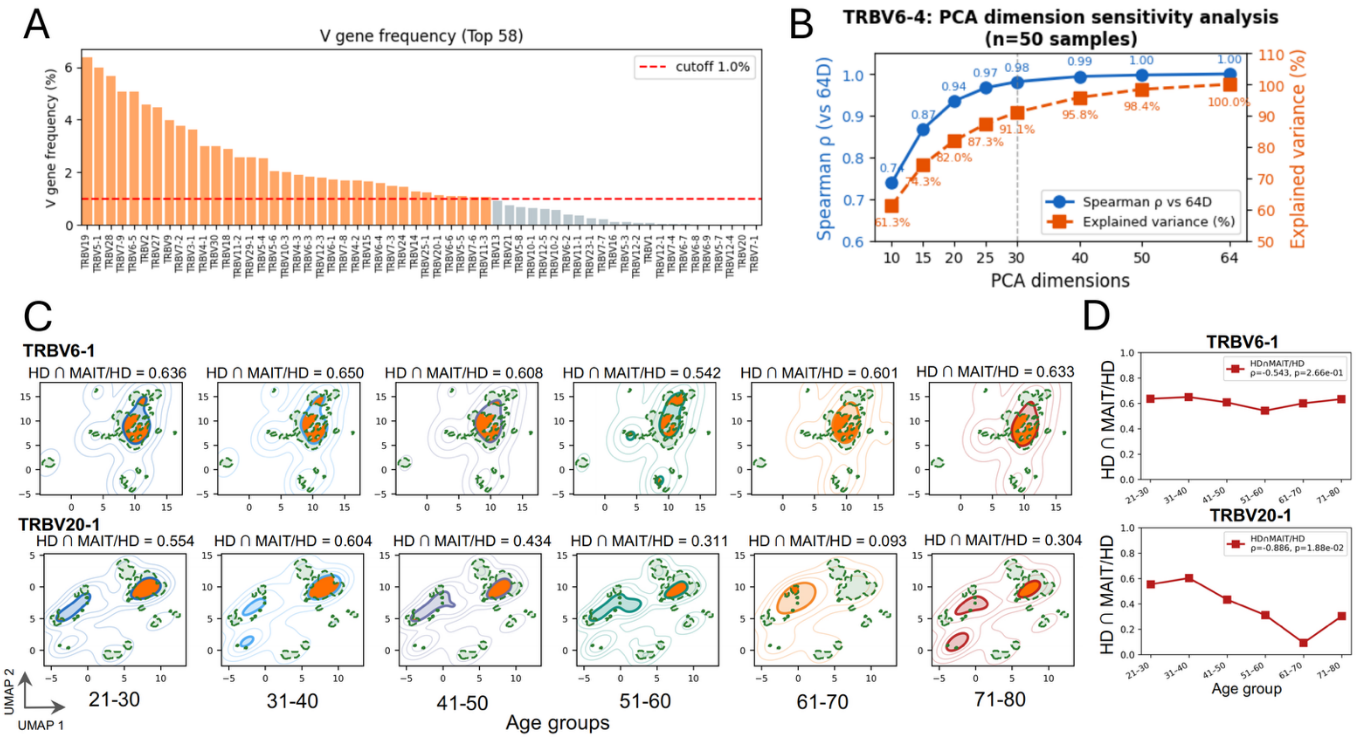


**Extended Data Fig. 5. Additional analyses of age-associated changes in the TCR repertoire landscape.**

**(A)** Frequencies of the 58 most abundant TRBV genes in the healthy reference repertoire. The dashed line indicates the 1% frequency cutoff. 35 TRBV genes above this cutoff are highlighted in orange and were selected for construction of the reference map. **(B)** Sensitivity analysis of the number of principal component analysis (PCA) dimensions used for TRBV6-4 repertoire mapping. For 50 samples, embeddings generated using different numbers of PCA dimensions were compared with the 64-dimensional embedding using Spearman correlation; cumulative explained variance is also shown. **(C)** Density distributions of TRBV6-1 and TRBV20-1 across age groups in the Gittelman cohort. TRBV6-1 and TRBV20-1 were selected as two additional V genes frequently used by mucosal-associated invariant T (MAIT) cells. For each age group, the high-density (HD) region was defined as the region above the 95th percentile of the density distribution for each age group. HD and MAIT regions are indicated by solid and dashed outlines, respectively, with their overlap highlighted. Values above each panel indicate the fraction of the HD region overlapping the MAIT region. **(D)** Fraction of the HD region overlapping the MAIT region across age groups for TRBV6-1 and TRBV20-1. Spearman correlations with age are shown.

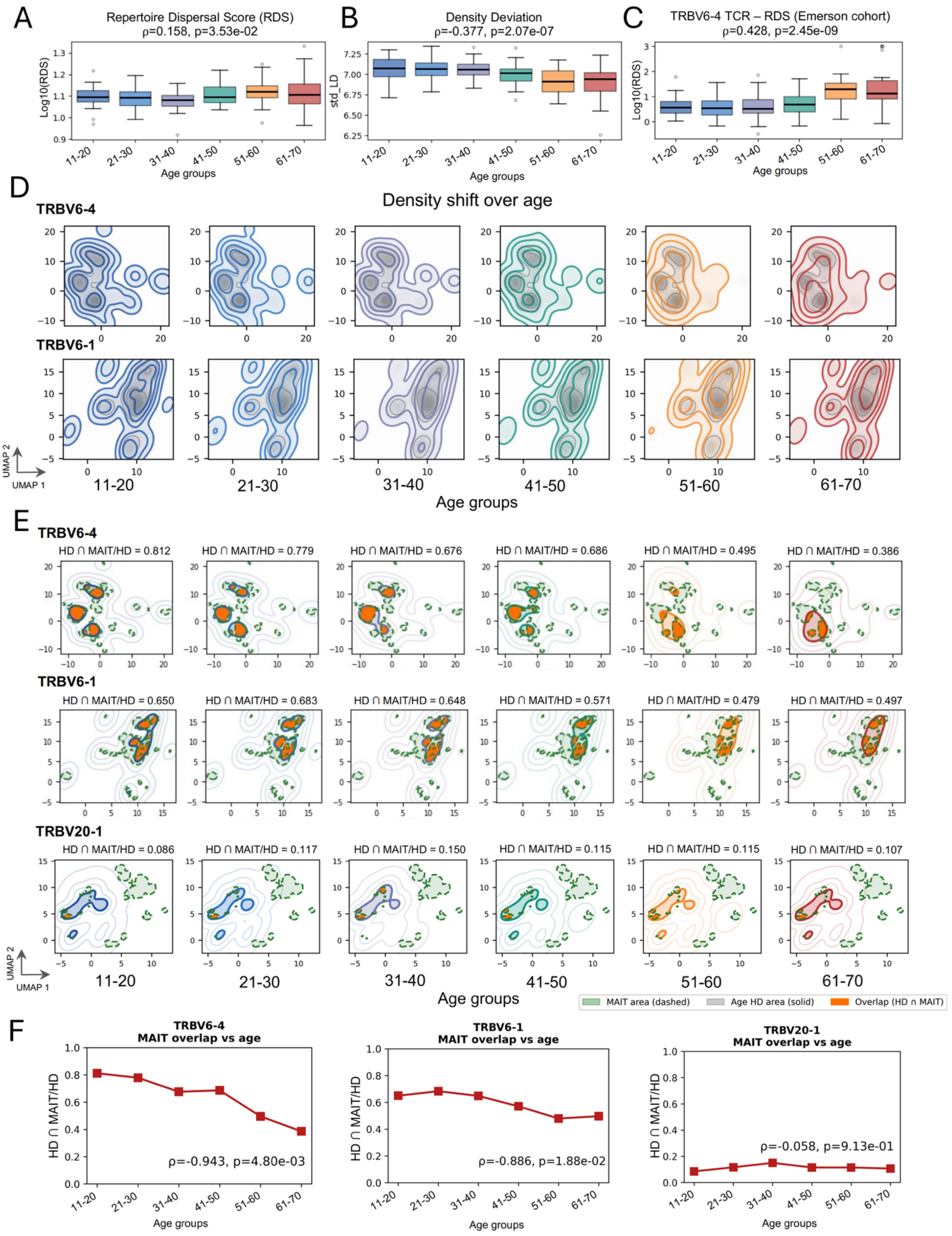
A
Repertoire Dispersal Score (RDS)
ρ=0.158, p=3.53e-02
Log10(RDS)
Age groups
B
Density Deviation
ρ=-0.377, p=2.07e-07
std_LD
Age groups
C
TRBV6-4 TCR – RDS (Emerson cohort)
ρ=0.428, p=2.45e-09
Log10(RDS)
Age groups
11-20
21-30
31-40
41-50
51-60
61-70
D
Density shift over age
TRBV6-4
TRBV6-1
UMAP 2
UMAP 1
Age groups
E
TRBV6-4
HD ∩ MAIT/HD = 0.812
HD ∩ MAIT/HD = 0.779
HD ∩ MAIT/HD = 0.676
HD ∩ MAIT/HD = 0.686
HD ∩ MAIT/HD = 0.495
HD ∩ MAIT/HD = 0.386
TRBV6-1
HD ∩ MAIT/HD = 0.650
HD ∩ MAIT/HD = 0.683
HD ∩ MAIT/HD = 0.648
HD ∩ MAIT/HD = 0.571
HD ∩ MAIT/HD = 0.479
HD ∩ MAIT/HD = 0.497
TRBV20-1
HD ∩ MAIT/HD = 0.086
HD ∩ MAIT/HD = 0.117
HD ∩ MAIT/HD = 0.150
HD ∩ MAIT/HD = 0.115
HD ∩ MAIT/HD = 0.115
HD ∩ MAIT/HD = 0.107
MAIT area (dashed)
Age HD area (solid)
Overlap (HD ∩ MAIT)
F
TRBV6-4
MAIT overlap vs age
ρ=-0.943, p=4.80e-03
TRBV6-1
MAIT overlap vs age
ρ=-0.886, p=1.88e-02
TRBV20-1
MAIT overlap vs age
ρ=-0.058, p=9.13e-01
HD ∩ MAIT/HD
Age groups

**Extended Data Fig. 6. Validation of age-associated repertoire changes in the Emerson cohort.**

**(A, B)** Overall **(A)** Repertoire Dispersal Score (RDS) and **(B)** density deviation across age groups in the Emerson cohort. Spearman correlations with age are shown. **(C)** TRBV6-4 RDS across age groups in the Emerson cohort. Spearman correlation with age is shown. **(D)** Density distributions of TRBV6-4 and TRBV6-1 across age groups. For each age group, the colored density shows the distribution of TCRs from that age group, while the gray background shows the distribution of the young healthy reference. **(E)** Density distributions of TRBV6-4, TRBV6-1, and TRBV20-1 showing their overlap with MAIT cell regions across age groups. For each age group, the high-density (HD) region was defined as the region above the 95th percentile of density within its corresponding map. HD and MAIT regions are indicated by solid and dashed outlines, respectively, with their overlap highlighted. Values above each panel indicate the fraction of the HD region overlapping the MAIT region. **(F)** Fraction of the HD region overlapping the MAIT region across age groups for TRBV6-4, TRBV6-1, and TRBV20-1. Spearman correlations with age are shown.

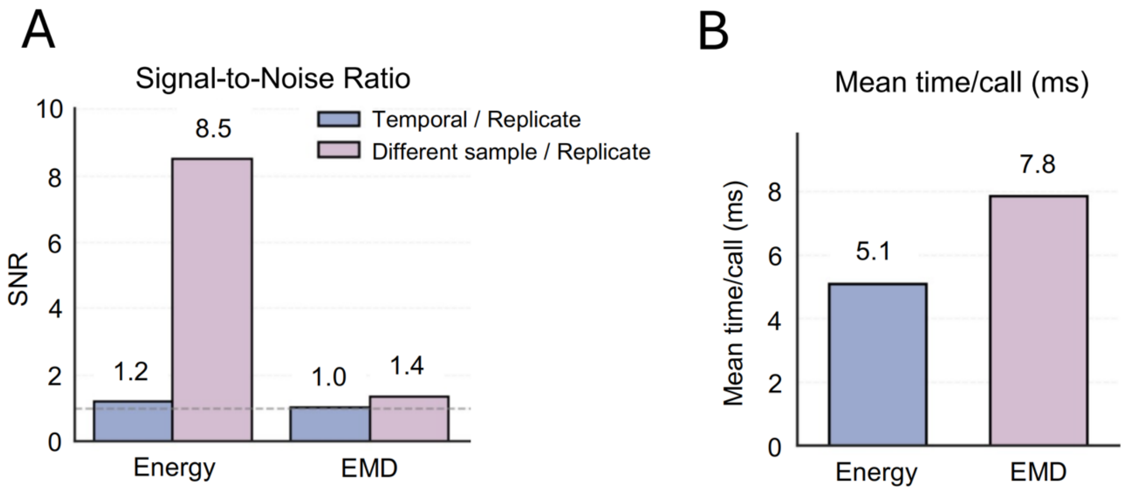


**Extended Data Fig. 7. Performance of Energy Shift for repertoire comparison.**

**(A)** Signal-to-noise ratio (SNR) of Energy Shift and EMD for temporal and different-sample comparisons relative to technical replicates. Technical replicates represent samples from the same individual at the same time point, whereas temporal pairs represent samples from the same individual at different time points. SNR was calculated as the mean distance between samples with biological differences divided by the mean distance between technical replicates. Higher SNR indicates greater separation of biological variation from technical-replicate variation. **(B)** Mean computation time per repertoire comparison for Energy Shift and Earth Mover's Distance (EMD).

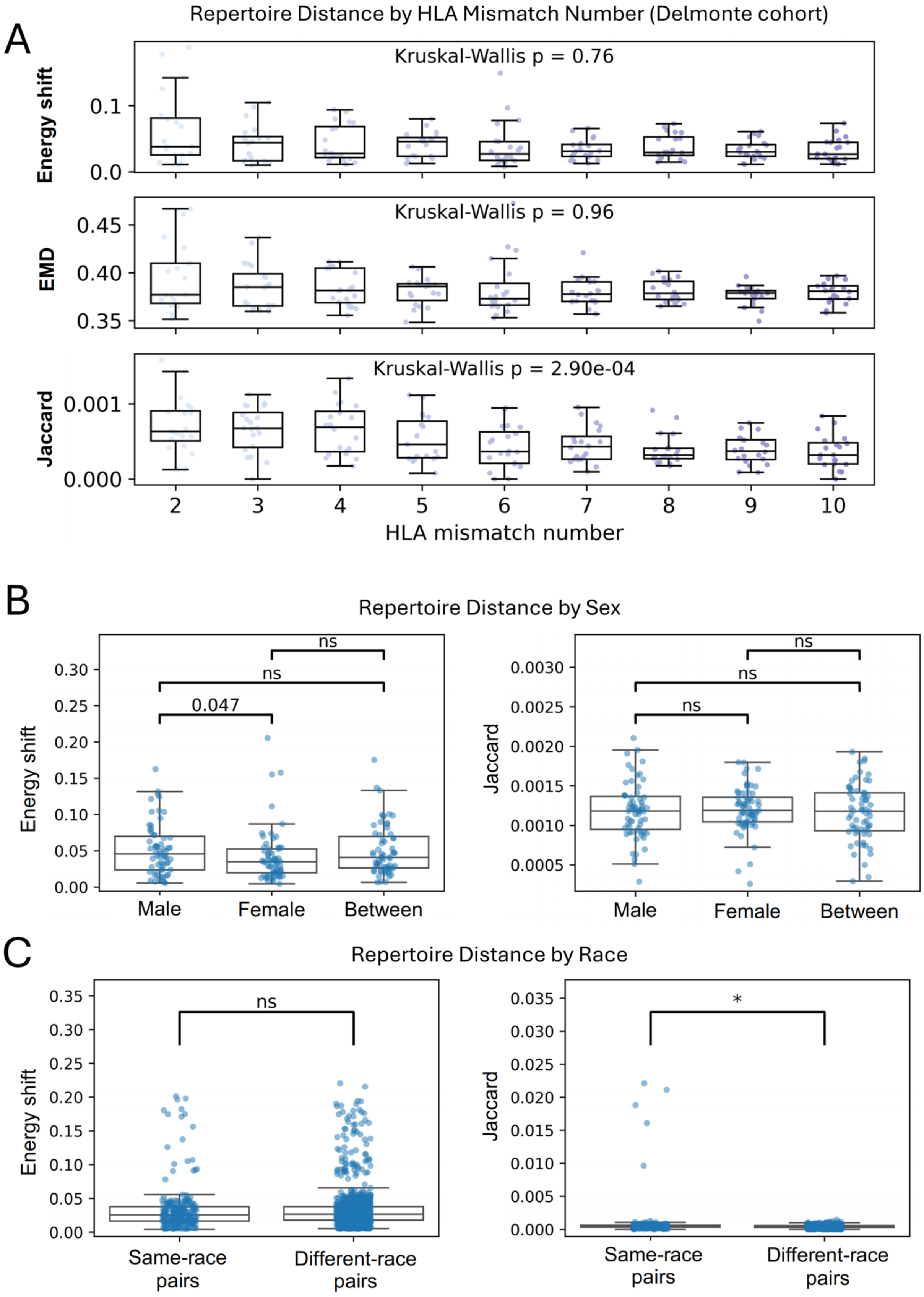


**Extended Data Fig. 8. Effects of HLA mismatch, sex, and race on repertoire distance.**

**(A)** Effect of human leukocyte antigen (HLA) mismatch on repertoire distance in the Delmonte cohort. Energy Shift, Earth Mover's Distance (EMD), and Jaccard index were calculated between sample pairs with different numbers of HLA mismatches. Kruskal-Wallis *P* values across HLA mismatch groups are shown. **(B)** Effect of sex on repertoire distance. Energy Shift and Jaccard index were compared among male-male, female-female, and male-female sample pairs. **(C)** Effect of race on repertoire distance. Energy Shift and Jaccard index were compared between same-race and different-race sample pairs. Statistical significance in **(B, C)** was assessed using two-sided Mann-Whitney U tests. ns, not significant; $P$ value < 0.05.

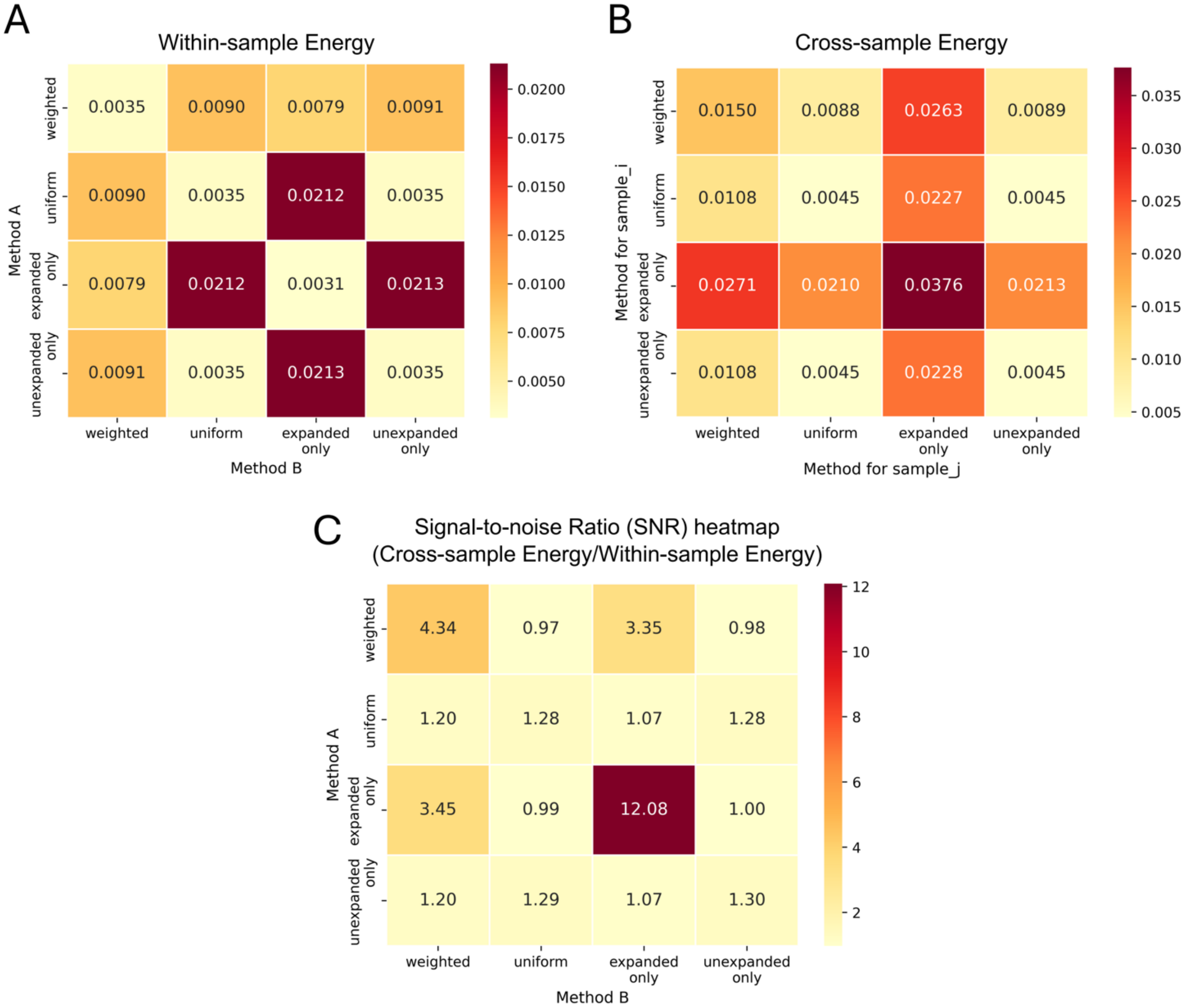


**Extended Data Fig. 9. Selection of repertoire sampling strategy for Energy Shift.**

**(A)** Mean within-sample Energy Shift for pairwise combinations of four repertoire sampling strategies: weighted, uniform, expanded-only, and unexpanded-only sampling. Weighted sampling sampled clonotypes according to their clone counts, whereas uniform sampling assigned equal sampling probability to all clonotypes. Expanded-only and unexpanded-only sampling used weighted sampling but were restricted to clonotypes with clone counts >1 and =1, respectively. Lower within-sample Energy Shift indicates less sampling-induced variation. **(B)** Mean cross-sample Energy Shift for pairwise combinations of the four sampling strategies defined in **(A)**. **(C)** Signal-to-noise ratio (SNR) for each combination of sampling strategies, calculated as the mean cross-sample Energy Shift divided by the mean within-sample Energy Shift. Higher SNR indicates greater between-repertoire differences relative to sampling-induced variation.
